# Compression injury regulates astrocyte morphology, metabolic function, and extracellular matrix modification in a 3D hydrogel

**DOI:** 10.1101/2025.06.20.660800

**Authors:** Ana N. Strat, Suhani Patel, Minh-Tri Ho Thanh, Alexander Kirschner, Souvik Ghosh, Michael P. Geiss, Arunkumar Venkatesan, Audrey M. Bernstein, Mariano Viapiano, Yutao Liu, Alison E. Patteson, Samuel Herberg, Preethi S. Ganapathy

## Abstract

In glaucoma, the optic nerve head (ONH) is exposed to increased biomechanical strain, impacting the resident astrocytes that maintain neural homeostasis. During disease progression astrocytes exhibit morphologic and metabolic shifts; however, the specific impact of glaucoma-related biomechanical strains on astrocyte behavior remains poorly understood. To address this, we used our previously established 3D cell-encapsulated extracellular matrix (ECM) hydrogel to investigate ONH astrocyte cellular and transcriptomic responses to varying biomechanical strain levels over time. Murine ONH astrocyte were encapsulated within an ECM hydrogel made from photocrosslinkable collagen type I and hyaluronic acid, and subjected to 0, 3, or 10% cyclic compression for 4h and 24h. We found significant restructuring of cytoskeletal morphology, metabolic dysregulation, and astrocyte-mediated ECM modulation that were strain-, duration- and hydrogel subregion-dependent. These phenotypic alterations were associated with diverse transcriptional changes in genes related to cell cycle and morphology, inflammation, metabolism and matrix remodeling that were driven by compressive strain intensity and duration. Our work reveals the direct role of compressive strain in eliciting a complex astrocyte response, supports targeting mechanosensation to prevent these pathologic astrocyte responses, and establishes ECM-based hydrogels as a platform to test mechanisms driving astrocyte mechanodysfunction. Altogether, our study offers new insights into astrocyte responses to biomechanical insult and demonstrates the use of a tunable 3D ECM hydrogel for future mechanistic studies of neurodegeneration.

**Graphical abstract:** 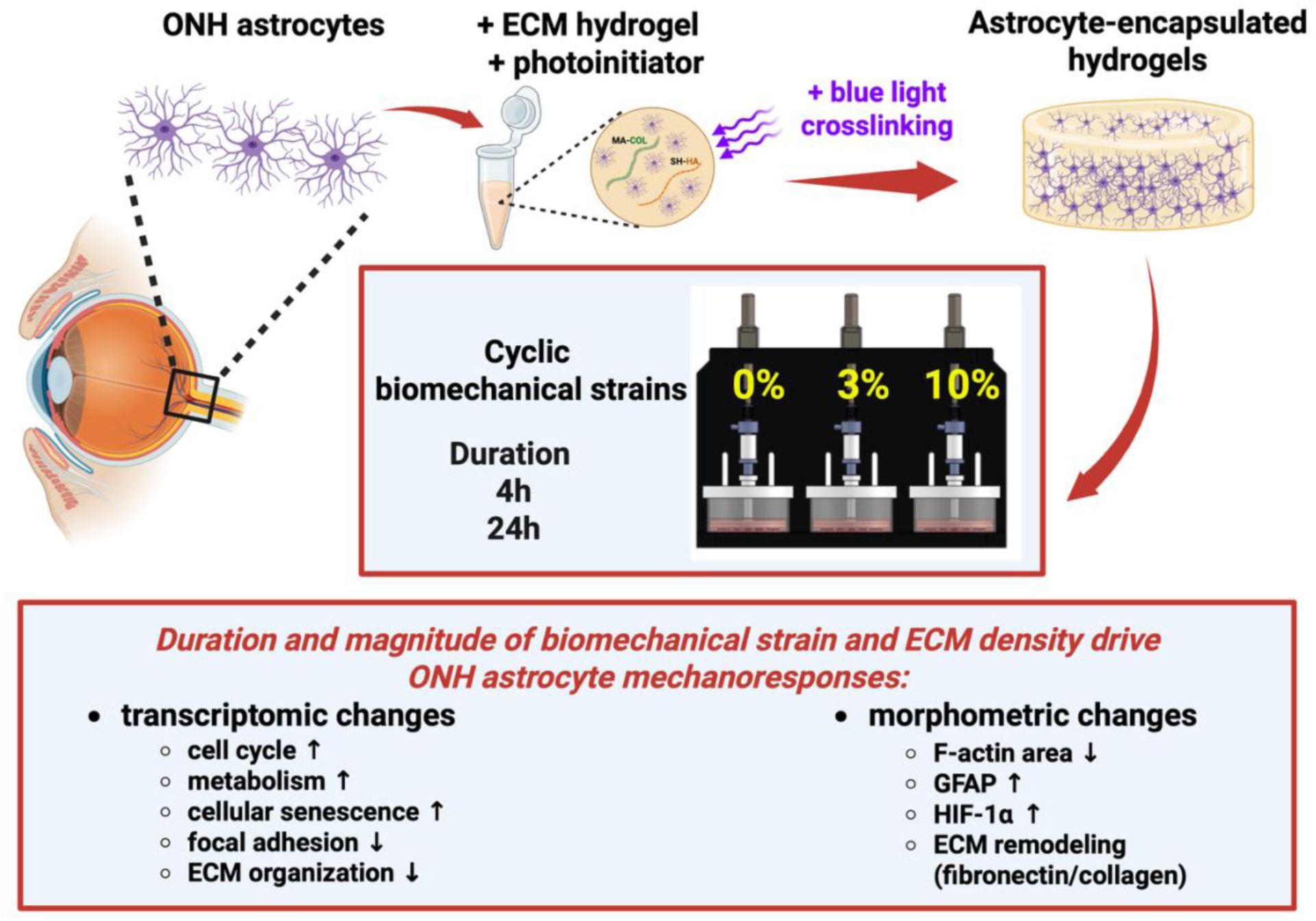

## 1. Introduction

Astrocytes are the master glial cells regulating central nervous system homeostasis. They provide trophic and matrix support to neurons, contribute to modulating synaptic activity and interact with neighboring glial cells and blood vessels to ensure optimal function and protection of neuronal tissue [1–3]. Astrocytes are highly sensitive to biomechanical stimuli, with their development and heterogeneity influenced by the surrounding biomechanical environment [4, 5]. Equally important to healthy physiologic conditions is the response of astrocytes to different biomechanical insults within a diseased central nervous system. In neuropathies induced acutely by trauma (e.g., traumatic brain or spinal cord injuries) or chronically by neurodegeneration (e.g., glaucoma), astrocytes are subjected to pathological biomechanical stimuli of various types, magnitudes, and durations that govern the cellular mechanoresponse [6, 7].

In glaucoma, pathologic biomechanical strains are directed upon the optic nerve head (ONH), where there is progressive loss of retinal ganglion cells ultimately leading to irreversible blindness [8, 9]. The local astrocytes within this region provide key support to axons and are first responders to acute glaucomatous insult prior to neuronal dysfunction [10, 11]. Their early response is characterized by a morphologic shift, associated with cytoskeletal protein alterations [10, 12–14]. As the disease progresses, astrocytes decouple from neighboring cells and remodel the surrounding extracellular matrix (ECM) by significantly increasing production of ECM proteins (e.g., fibronectin and collagen IV), matrix crosslinking/degrading enzymes (e.g., transglutaminase 2 and matrix metalloproteinases 2/9), and pro-fibrotic cytokines (e.g., transforming growth factor beta 2 (TGFβ2)) [15, 16]. These glial cells are also responsible for matching the metabolic demand of neurons under stress. This metabolic interdependence, exemplified by the glutamate–glutamine cycle and astrocyte-mediated vasodilation, is crucial for neuronal health and its disruption contributes to neurodegenerative diseases [3, 17].

Evidence supports that the ONH in glaucoma is subjected to various degrees of compressive and tensile strains that vary by microregion. Specifically, finite element modeling studies predict between 3-10% compressive and 2-5% tensile biomechanical strain on the ONH with intraocular pressure (IOP) elevation [18–20]. While many studies have used such modeling approaches to understand the biomechanical strains within the glaucomatous ONH, there are limited reports on the direct phenotypic and transcriptional responses of local astrocytes to isolated biomechanical strains. This is a crucial gap in our understanding of how glaucomatous biomechanical strains can impact astrocyte behavior and, ultimately, disrupt their functional support of neurons. Prior studies have reported on ONH astrocyte cell cultures subjected to mechanical stressors such as hydrostatic pressure and tension in 2D [17, 21]. While not without merit, conventional 2D *in vitro* culture systems are not suitable to analyze 3D astrocyte morphological complexity and coupling under biomechanical forces. Furthermore, it is nearly impossible to subject monolayer astrocyte cultures to compression in 2D.

We previously developed a cell–encapsulated hydrogel containing murine ONH astrocytes and ECM biopolymers (collagen type I and hyaluronic acid), which permits detailed analysis of cell shape and cell–cell/cell–ECM interactions in 3D [22]. In the present study, we investigated how ONH astrocytes that are encapsulated within this ECM hydrogel respond to varying degrees of compressive biomechanical strain over time using a well-established bioreactor [23–25]. We aimed to thoroughly characterize the phenotypic and transcriptional responses of astrocytes to strain magnitude and duration. To achieve this, we conducted histologic analyses and bulk RNA sequencing to determine changes in cytoskeletal organization, metabolic function, ECM remodeling, and gene expression over time. Ultimately, our study represents a comprehensive investigation of time–dependent ONH astrocyte mechanoresponses to glaucoma–relevant biomechanical strains and seeks to provide mechanistic insights into broader astrocyte mechanodysfunction in neurodegenerative diseases of the central nervous system.

## 2. Results

### 2.1 Characterization of astrocyte-encapsulated hydrogel pore size, permeability, and regional differences

Under healthy conditions, local astrocytes create a honeycomb network spanning the diameter of the rodent ONH [26]. Their processes can measure a thickness of 1.3 ± 0.04 μm with elongations of up to 30 μm [27]. To determine whether our previously developed ECM hydrogel was adequately porous to allow for mouse ONH astrocyte (MONHA) process elongation and coupling through the collagen network, we first quantified the pore size of acellular ECM hydrogels in the absence of biomechanical strain. Using confocal reflectance microscopy to directly visualize collagen fibers within acellular hydrogels, we observed that the collagen fibrils displayed a homogeneous network with pore sizes ranging from 2 – 7 μm (Suppl. Fig. 1A-B).

Next, we quantified ECM hydrogel permeability over time (Suppl. Fig. 1C). Since our astrocyte growth media contains primarily fetal bovine serum (FBS) proteins ranging < 60 – 70 kDa (with albumin being the most abundant) [28], and added epidermal growth factor which is ∼ 6 kDa [29], we chose to use a 70 kDa FITC-labelled dextran tracer solution as an indicator for permeability. Both the acellular and MONHA-encapsulated hydrogels showed robust permeability, as seen by the rapid detection of increased fluorescence below the ECM hydrogel within 5 minutes of adding FITC-labelled dextran solution to the top of the hydrogels (Suppl. Fig. 1C). Thus, these results support that MONHA-encapsulated hydrogels allow for efficient and adequate nutrient distribution.

**Supplemental Fig. 1.**
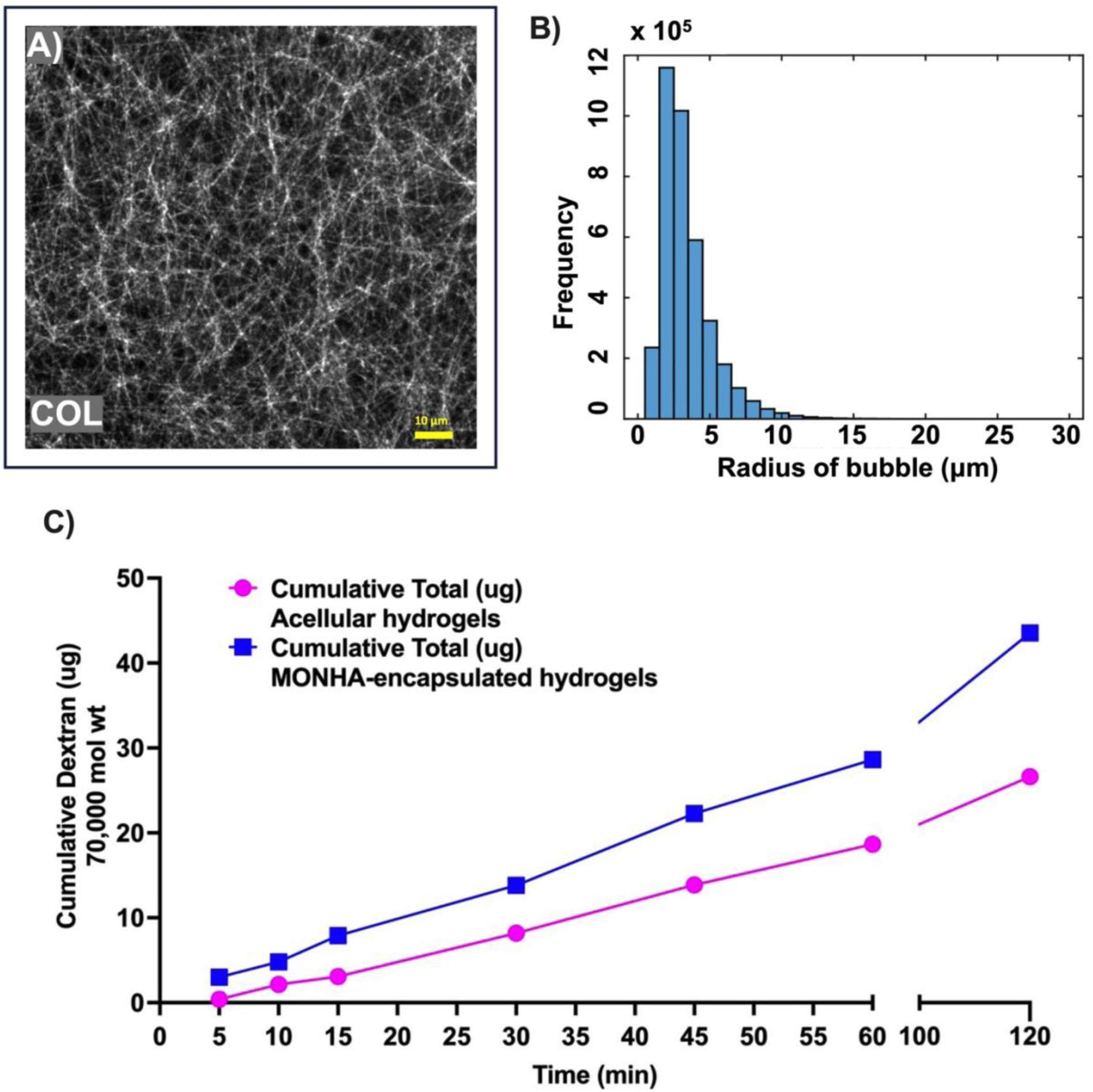
Characterization of the MONHA-encapsulated ECM hydrogel model. **(A)** Representative fluorescence image of collagen fibril distribution and **(B)** quantification of pore size within acellular hydrogels (N = 5 hydrogels). Scale bar 10 μm. **(C)** Assessment of the normalized average permeability of acellular and MONHA-encapsulated hydrogels (N = 3 hydrogels/group) as calculated by FITC-labelled 70kDa dextran as tracer molecule.

To characterize the architecture of the ECM network within our MONHA-encapsulated hydrogels, we evaluated the compaction of collagen fibrils. We observed regional differences within our constructs such that samples at baseline (under no biomechanical strain) displayed increased collagen compaction in the periphery of the ECM hydrogel as compared to the core (Suppl. Fig 2A-B). Moreover, the collagen compaction appeared greater immediately adjacent to astrocyte cell bodies/processes (Suppl. Fig. 2A). Collectively, these data support that encapsulated ONH astrocytes differentially remodel and contract their surrounding collagen fibers in the core as compared to the unbound periphery of the hydrogel. Because of this heterogeneity in collagen compaction, we artificially delineated the core from the periphery of MONHA-encapsulated hydrogels for subsequent experiments and comparisons (Suppl. Fig. 2C).

**Supplemental Fig. 2.**
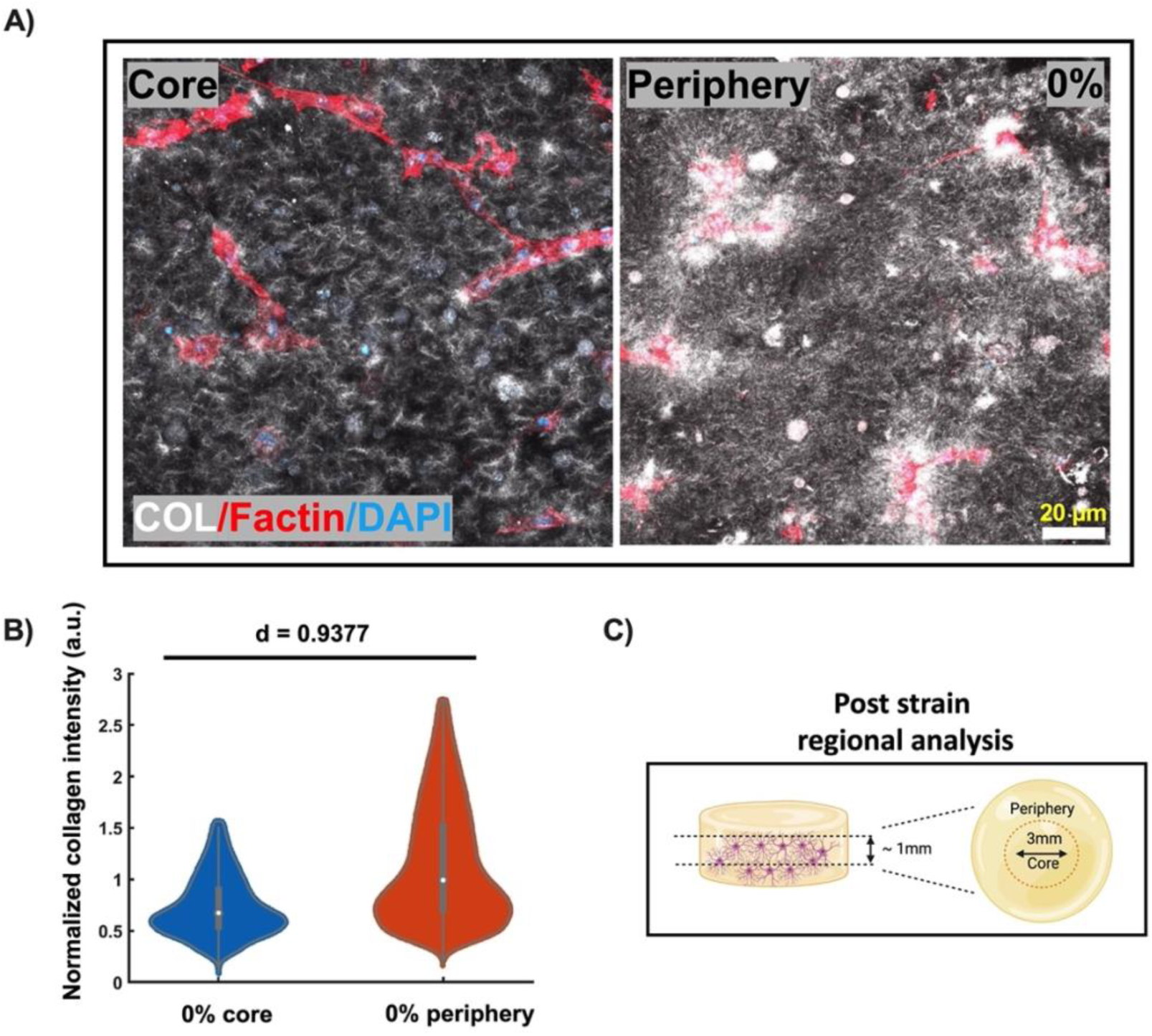
Regional differences in collagen fibril compaction of MONHA-encapsulated hydrogels. **(A)** Density of collagen fibrils in the core and periphery of the MONHA-encapsulated hydrogels after 2 weeks in culture; scale bar 20 μm. **(B)** Quantitative analysis of normalized collagen compaction within the core and periphery of MONHA-encapsulated hydrogels (N = 3 hydrogels/group). Cohen’s d = 0.20, 0.50, and 0.80 is used to indicate the effect sizes as small, medium, or large, respectively. **(C)** Schematic delineating regional segmentation of MONHA-encapsulated hydrogels post biomechanical strain.

### 2.2 Quantification of astrocyte cell viability within MONHA-encapsulated hydrogels after compressive strain

After characterizing the ECM hydrogel, we then applied compressive biomechanical strains to MONHA-encapsulated hydrogels using a custom bioreactor [23–25]. We used 3% and 10% compressive strains at a frequency of 1 Hz, which have been reported in ONH tissues during glaucomatous insult [18–20]. After 24h compression, we quantified astrocyte cell viability in MONHA-encapsulated hydrogels (Suppl. Fig. 3). We found no detectable difference in total viability by MTS assay after compression for 24h (Suppl. Fig. 3).

**Supplemental Fig. 3.**
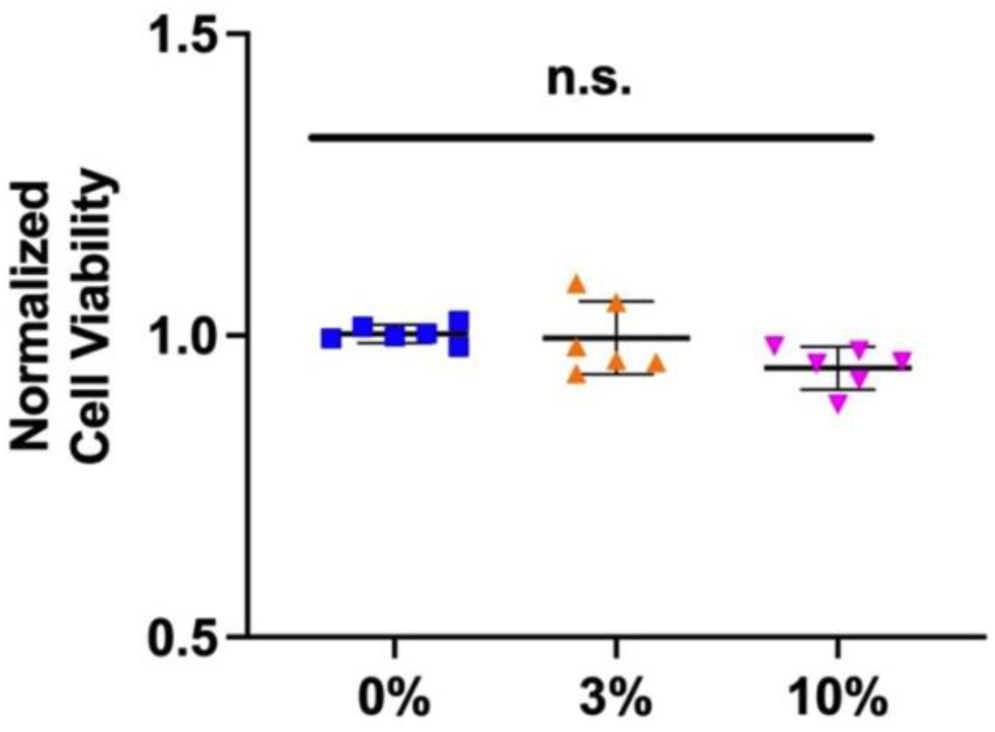
Cell viability following compressive strain. Normalized MONHA cell viability after 24h compression via MTS assay (N = 6 hydrogels/group). Significance was determined by one-way ANOVA using multiple comparisons tests.

### 2.3 Compressive strain alters MONHA cytoskeletal architecture

Immediately following glaucomatous insult, ONH astrocytes *in situ* respond by reorienting their cell bodies and remodeling their F-actin cytoskeleton architecture [12–14, 27]. Thus, we asked whether isolated compressive strains would induce similar F-actin and cytoskeletal reorganization. To do so, we quantified F-actin area coverage and levels of GFAP expression in MONHA-encapsulated hydrogels after biomechanical strain application for 4h and 24h, respectively.

After 4h compression, MONHAs showed no significant changes in F-actin area coverage in either the core or periphery of the ECM hydrogels (Fig. 1A-C). In contrast, after 24h compression, there was a significant drop in F-actin area coverage within the core of the hydrogel in a strain-dependent manner (Fig. 1A-C). Both 3% and 10% groups showed ∼50% decrease in F-actin area coverage compared to control (0%) (Fig. 1C), which was also associated with increased cell alignment in the 10% compression group (Suppl. Fig. 4A,B). Interestingly, application of compressive strain did not alter the F-actin area coverage or cell alignment in the periphery of the ECM hydrogel in either strain group (Fig. 1C; Suppl. Fig. 4C,D). Overall, these data demonstrate that encapsulated MONHAs are robustly mechanoresponsive, altering cellular morphology in response to biomechanical stimuli.

**Fig. 1.**
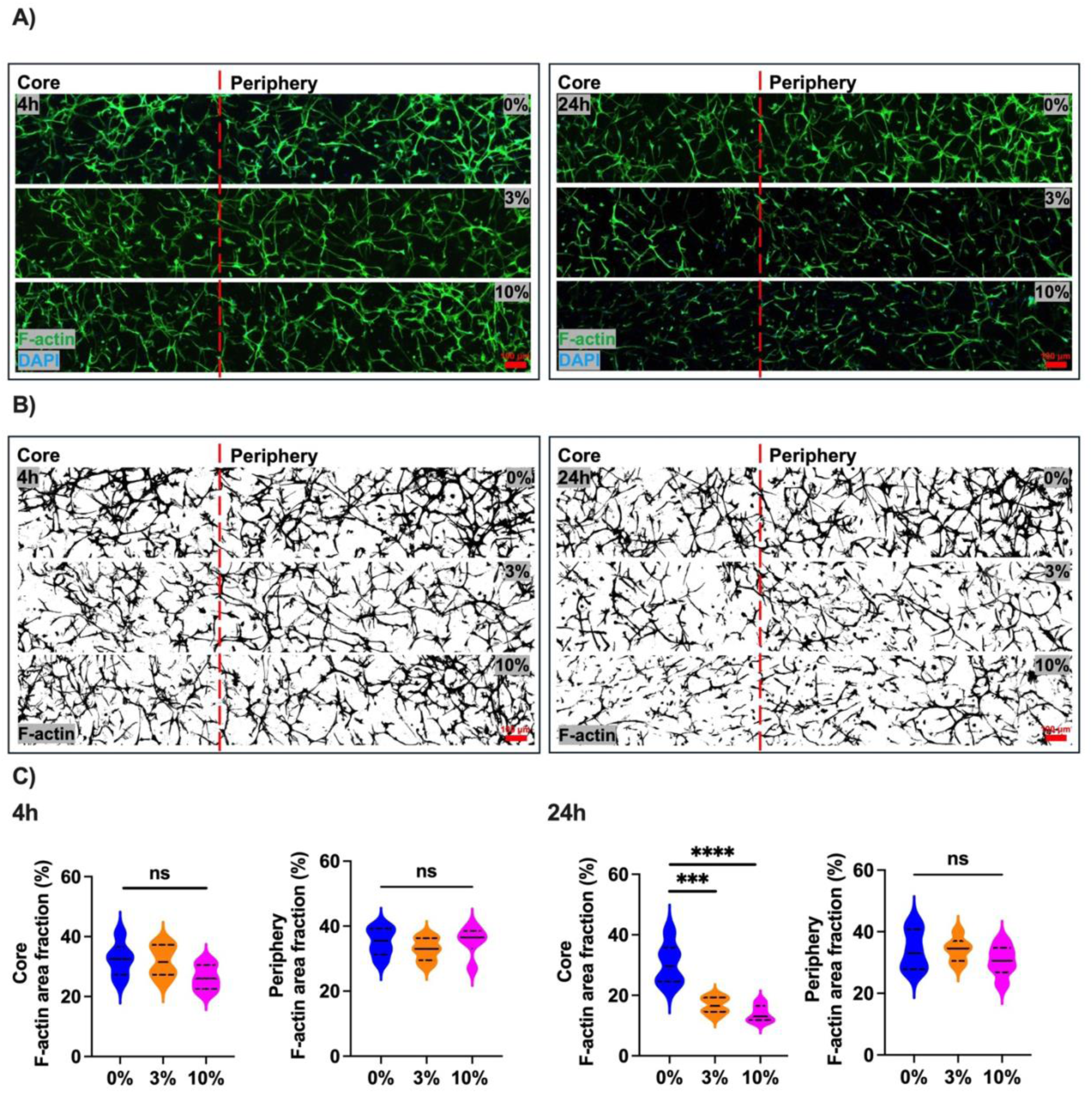
Regional differences in MONHA morphology in response to duration and magnitude of compressive strain. **(A)** Representative fluorescence images of astrocyte F-actin staining (A) and cytoskeletal binarized masks (B). Scale bar: 100 μm. **(C)** Quantitative analyses of F-actin mask area (n = 6 fields of view/group collected from N = 2-3 hydrogels/group). Statistical significance was determined using one-way ANOVA (*p < 0.05, **p < 0.01, ***p < 0.001, **** p < 0.0001) for F-actin area fraction analyses.

**Supplemental Fig. 4.**
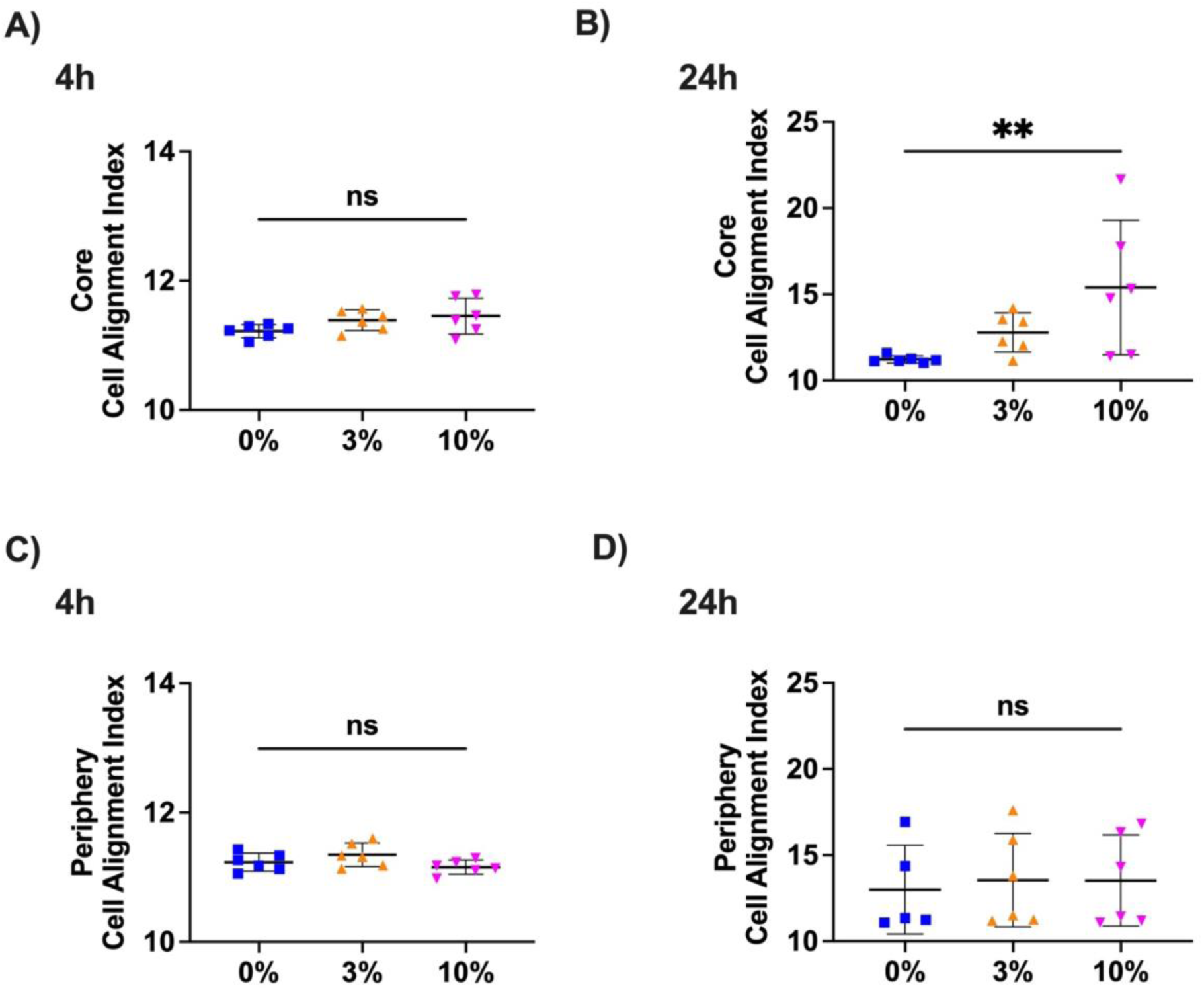
Regional differences in encapsulated MONHA cell alignment in response to compressive strain. Quantitative analyses of cell alignment based on F-actin mask area (n = 5-6 fields of view/group collected from N = 3 hydrogels/group) in response to 4h **(A)** and 24h of increasing biomechanical strain (i.e., 3% or 10%) **(B)** within the core of the hydrogel, and 4h **(C)** and 24h of biomechanical strain **(D)** within the periphery of the hydrogel, Statistical significance was determined using one-way ANOVA (** p < 0.01).

Astrocytes respond to glaucomatous insults within the ONH by undergoing reactive gliosis, in part characterized by an increase in expression of intermediate filaments such as GFAP [30, 31]. We previously showed that both MONHAs in 2D and encapsulated within 3D ECM hydrogels can be induced to express higher levels of GFAP upon stimulation with a known inducer of astrocyte reactivity such as exogeneous TGFβ2 [22, 32]. Therefore, we asked whether application of compressive strain would induce a similar induction in intermediate filaments. Control (0% compression) MONHA-encapsulated hydrogels showed baseline immunoreactivity of GFAP throughout the hydrogel. After both 4h and 24h compression, GFAP immunolabeling increased ∼1.5-fold in both the core and periphery of the ECM hydrogel (Fig. 2A-C, Suppl. Fig. 5A-C). As such, these data support that compressive strains directly induce cytoskeletal alterations in encapsulated astrocytes.

**Fig. 2.**
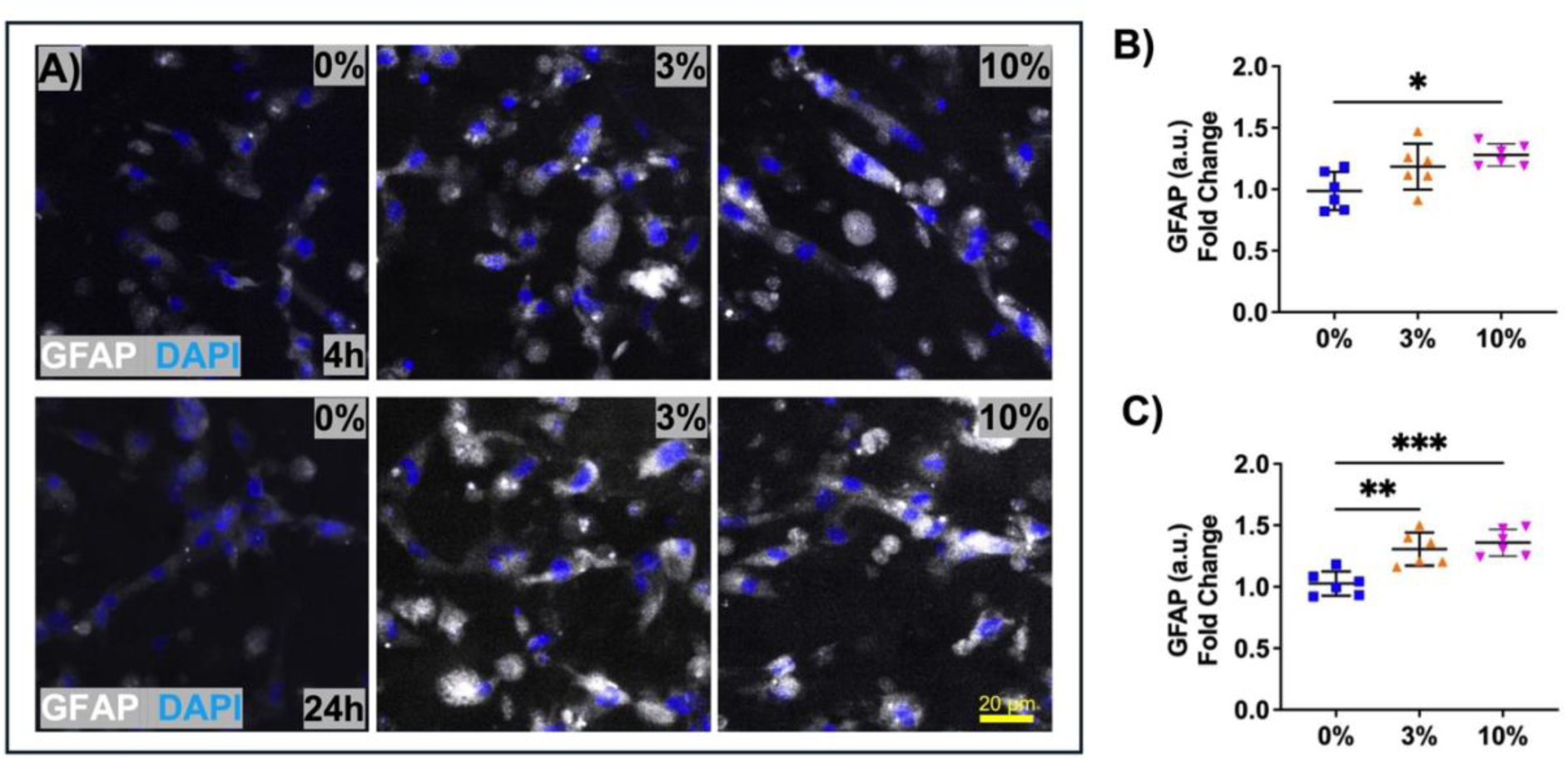
GFAP expression of encapsulated-MONHAs within the hydrogel core in response to compressive strain. **(A)** Representative fluorescence images and quantification of fold change in astrocyte GFAP signal intensity (n = 6 fields of view/group collected from N = 3 hydrogels/group) after **(B)** 4h and **(C)** 24h of increasing strain (i.e., 3% or 10%). Statistical significance was determined using one-way ANOVA (*p < 0.05, **p < 0.01, ***p < 0.001) for GFAP expression levels. Scale bar: 20 μm.

**Supplemental Fig. 5.**
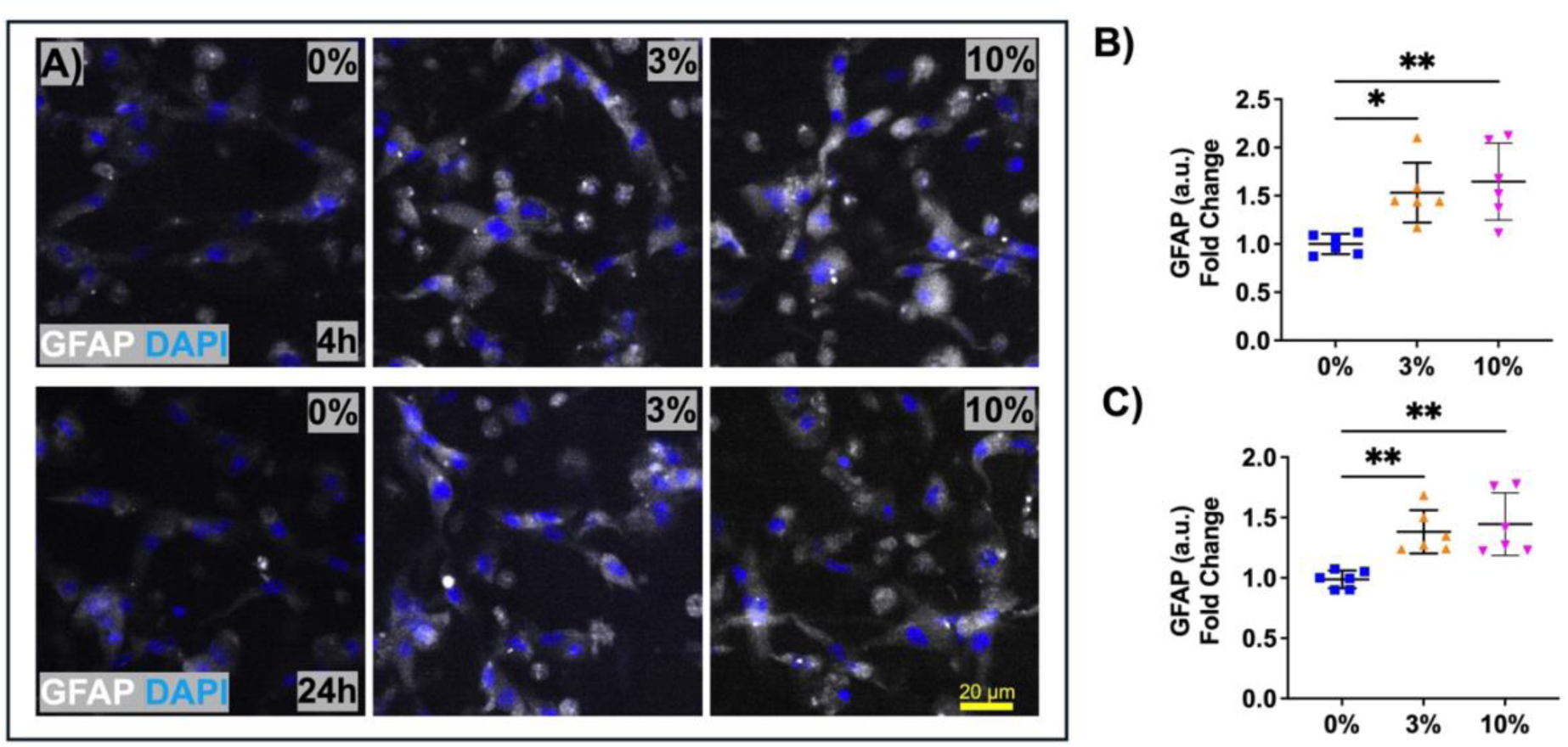
GFAP expression of encapsulated-MONHAs within the hydrogel periphery in response to compressive strain. **(A)** Representative fluorescence images and quantification of fold change in astrocyte GFAP signal intensity (n = 6 fields of view/group collected from N = 3 hydrogels/group) after **(B)** 4h and **(C)** 24h of increasing strain (i.e., 3% or 10%). Statistical significance was determined using one-way ANOVA (*p < 0.05, **p < 0.01) for GFAP expression levels. Scale bar: 20 μm.

### 2.4 Compressive strain alters HIF-1*α* levels within MONHA-encapsulated hydrogels

Glaucomatous insults are associated with hypoxia and metabolic stress within the retina and ONH since IOP-induced deformations can impact the local vasculature and reduce tissue oxygenation [33, 34]. Thus, we asked if astrocytes within our model could modulate expression of hypoxia-inducible factor (HIF), thereby potentially affecting downstream metabolic activity. Specifically, to determine whether encapsulated MONHAs undergo such metabolic stress in response to biomechanical strain, we examined levels of HIF-1α after application of compressive strain. HIF-1α immunoreactivity was low in control (0%) samples at both timepoints (i.e., 4h and 24h; Fig. 3, Suppl. Fig. 6) regardless of ECM hydrogel region (i.e., core vs periphery). Hydrogel-encapsulated MONHAs exhibited significantly increased HIF-1α levels by ∼1.25-fold in the core groups after 4h compression and by ∼2-fold in the periphery (Fig. 3A, B); HIF-1α also increased ∼1.5-2-fold after 24h compression (Fig. 3A, C). Similar trends of HIF-1α elevation were observed in the periphery of the ECM hydrogel (Suppl. Fig. 6). Taken together, these data support that compressive strain can directly induce changes in metabolic regulatory factors.

**Fig. 3.**
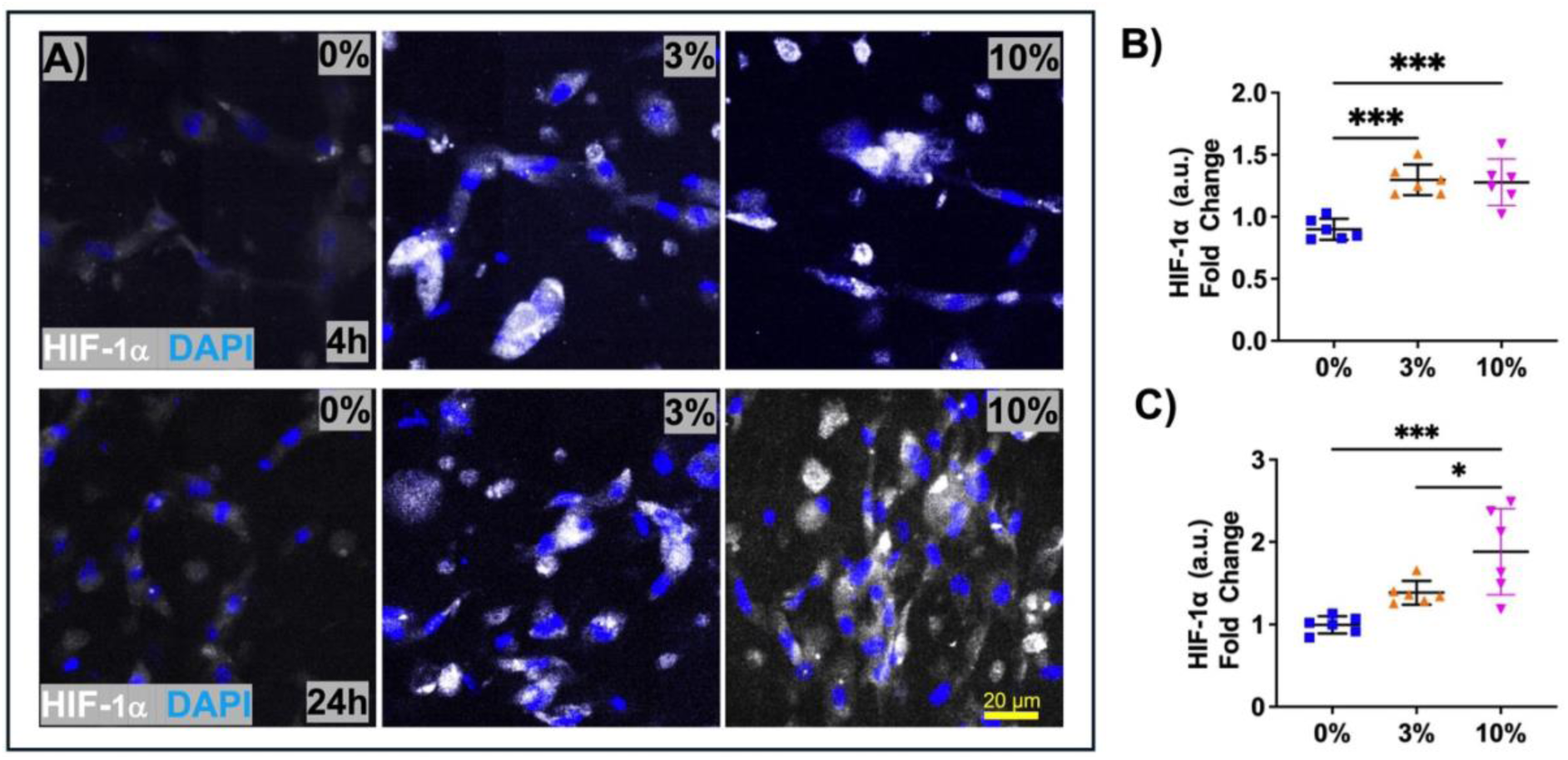
HIF-1α expression of encapsulated-MONHAs within the hydrogel core in response to compressive strain. **(A)** Representative fluorescence images and quantification of fold change in astrocyte HIF-1α expression signal intensity (n = 6 fields of view/group collected from N = 3 hydrogels/group) after **(B)** 4h and **(C)** 24h of increasing strain (i.e., 3% or 10%). Statistical significance was determined using one-way ANOVA (*p < 0.05, **p < 0.01, ***p < 0.001) for HIF-1α expression levels. Scale bar: 20 μm.

**Supplemental Fig. 6.**
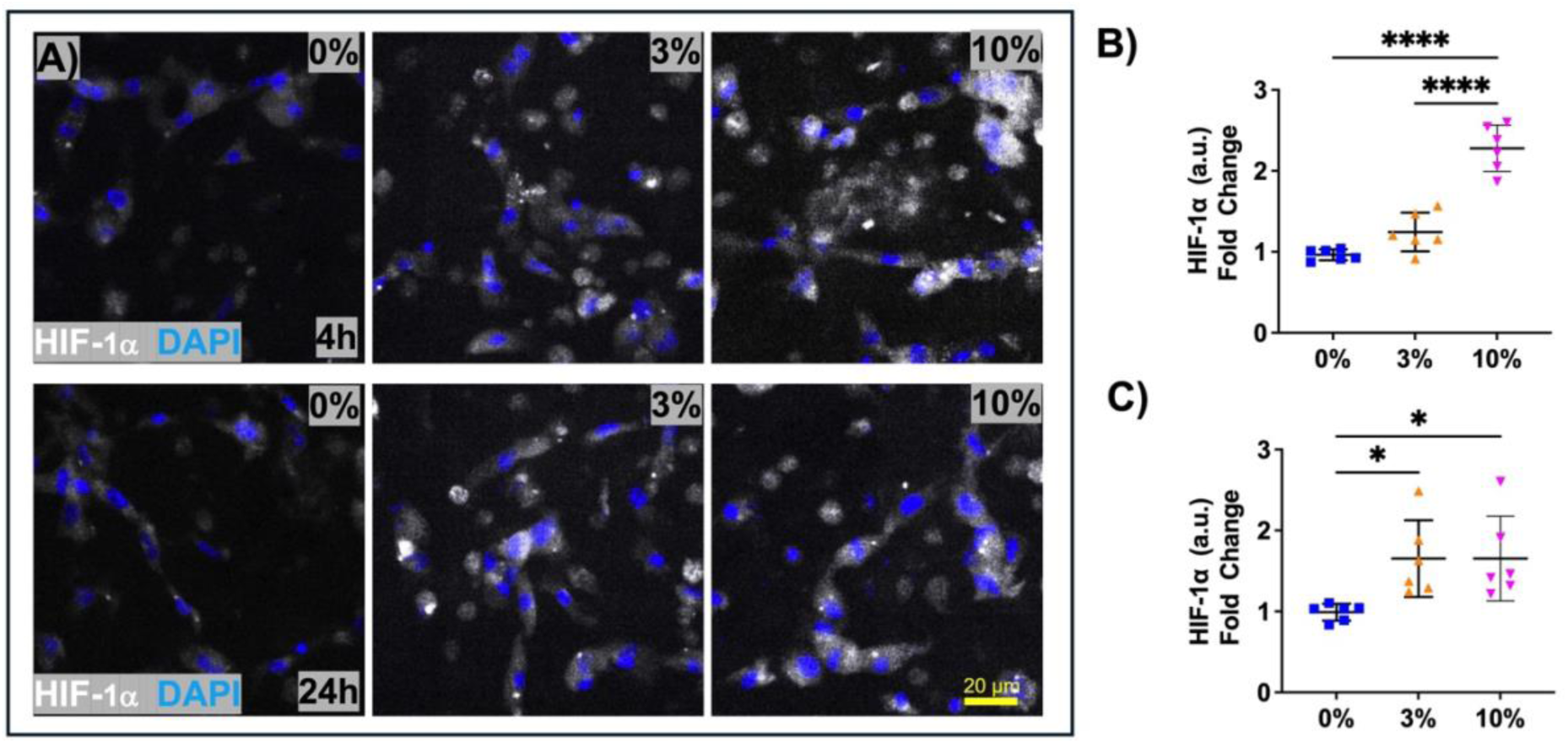
HIF-1α expression of encapsulated-MONHAs within the hydrogel periphery in response to compressive strain. **(A)** Representative fluorescence images and quantification of fold change in astrocyte HIF-1α expression signal intensity (n = 6 fields of view/group collected from N = 3 hydrogels/group) after **(B)** 4h and **(C)** 24h of increasing strain (i.e., 3% or 10%). Statistical significance was determined using one-way ANOVA (*p < 0.05, **** p < 0.0001) for HIF-1α expression levels. Scale bar: 20 μm.

### 2.5 Compressive strain on MONHA-encapsulated hydrogels induces matrix remodeling via collagen network remodeling and fibronectin deposition

*In situ,* ONH astrocytes modulate their surrounding ECM via secretion of crosslinking and/or degrading enzymes [15, 30]. Therefore, we asked how application of compressive strain would influence astrocyte modulation of the ECM microenvironment within our hydrogels. We first performed confocal reflectance microscopy to directly visualize collagen fibers within MONHA-encapsulated hydrogels after compression. We previously showed that control constructs (0%) display regional differences in collagen arrangement, with more uniformly distributed collagen fibers within the core of the ECM hydrogel, and an increased density of collagen fibers in the periphery of the hydrogel (Suppl. Fig. 2). While 4h compression did not induce significant remodeling of the collagen fibrils across groups, after 24h compression, both 3% and 10% showed significant *decrease* of collagen compaction within the core (Fig. 4A, B) and a significant *increase* in collagen compaction in the periphery compared to control (0%) (Suppl. Fig. 7A, B). Collectively, these data support that application of compressive strain for 24h significantly alters collagen compaction within ECM hydrogels in a region-dependent manner.

**Fig. 4.**
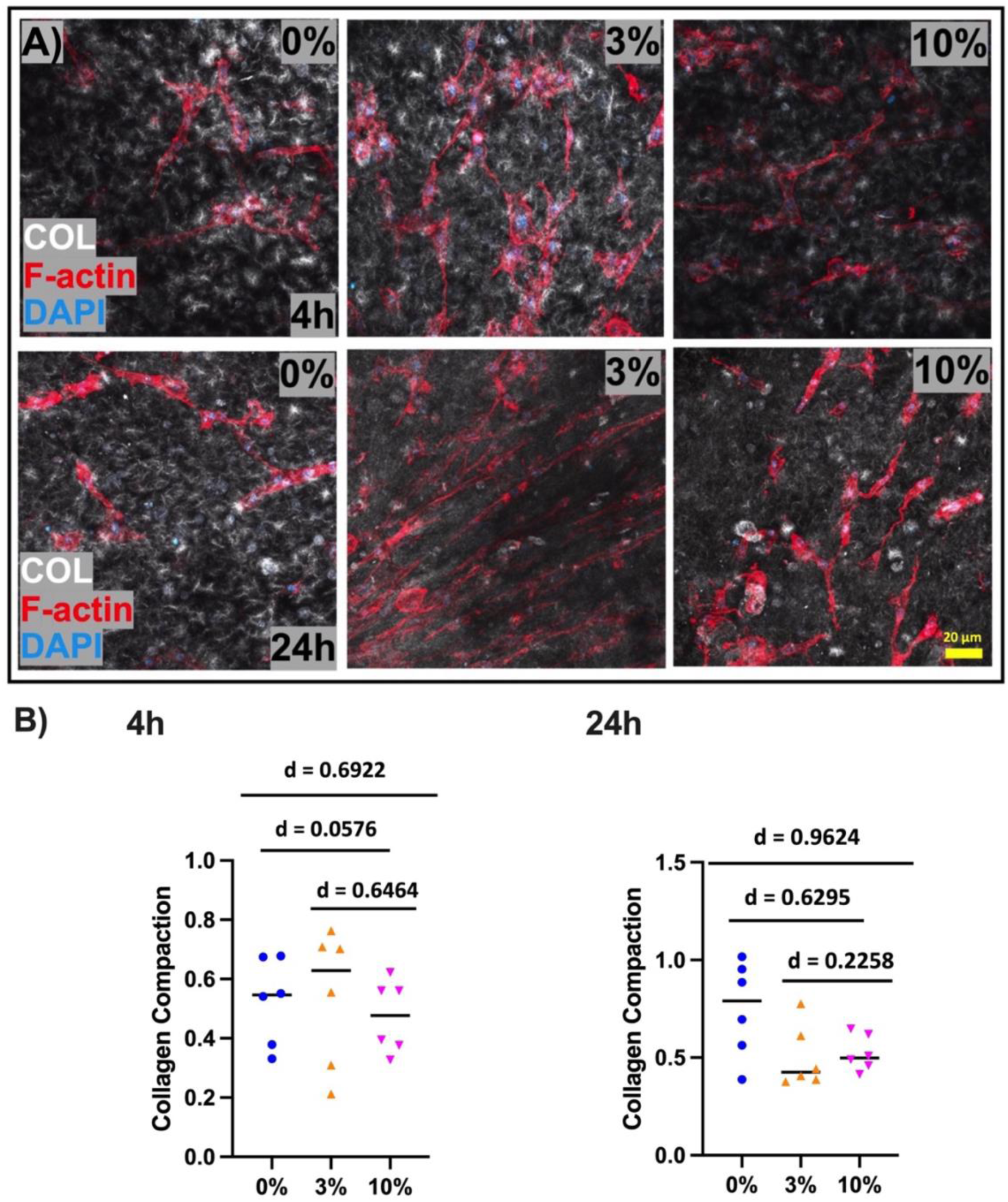
ECM collagen fibrils remodeling in the core of MONHA-encapsulated hydrogels in response to compressive strain. **(A)** Representative fluorescence images of collagen fibrils remodeling in response to duration (i.e., 4h or 24h) and magnitude of strain (i.e., 3% or 10%) (n = 6 fields of view/group collected from N = 3 hydrogels/group). Scale bar: 20 μm. **(B)** Quantification of collagen compaction after compression. Cohen’s effect size d (Cohen’s d = 0.20, 0.50, and 0.80 is used to indicate the effect sizes as small, medium, or large, respectively).

**Supplemental Fig. 7.**
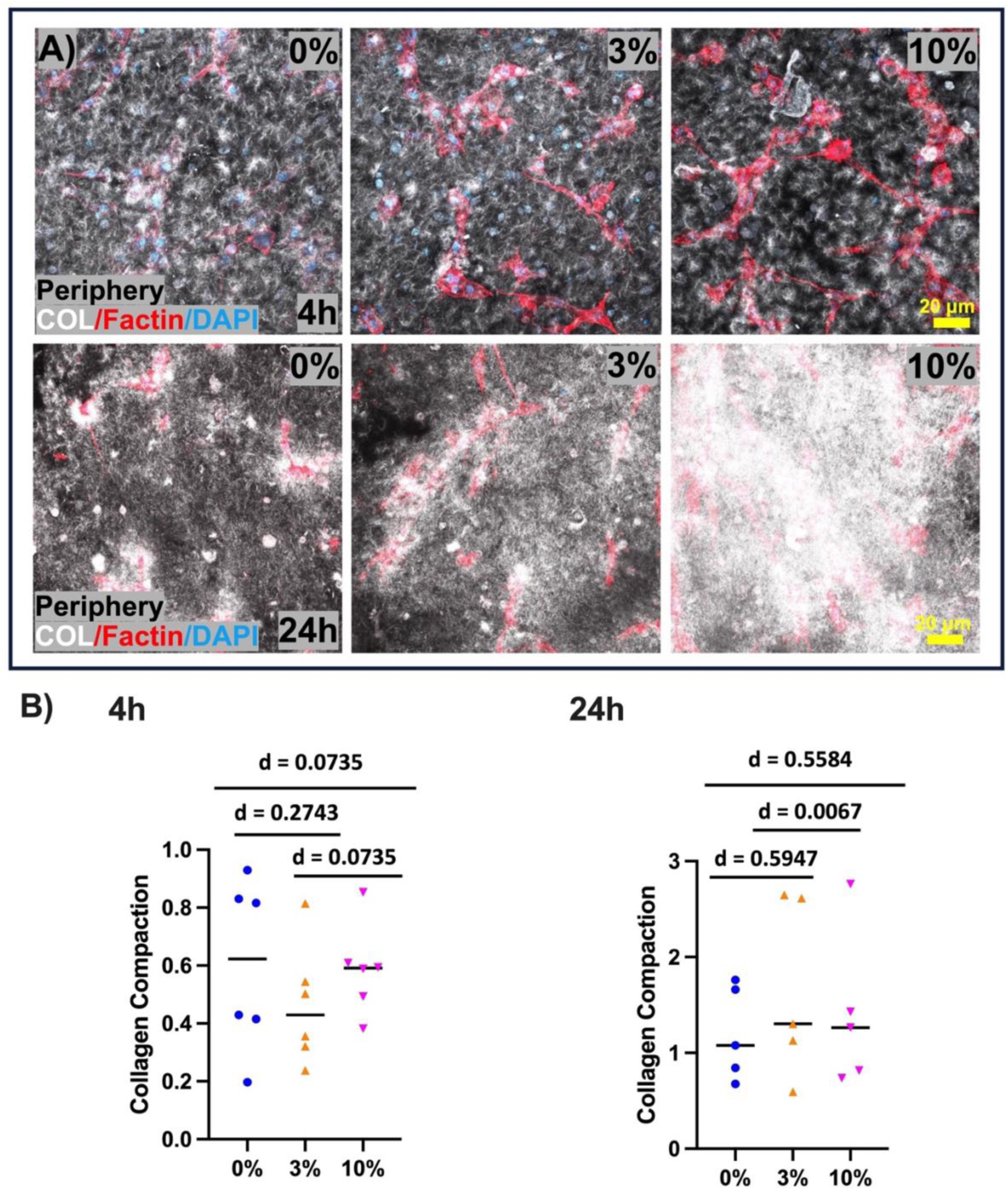
ECM collagen fibrils remodeling in the periphery of MONHA-encapsulated hydrogels in response to compressive strain. **(A)** Representative fluorescence images of collagen fibrils remodeling in response to duration (i.e., 4h or 24h) and magnitude of strain (i.e., 3% or 10%) (n = 5-6 fields of view/group collected from N = 3 hydrogels/group). Scale bar: 20 μm. **(B)** Quantification of collagen compaction after compression. Cohen’s effect size d (Cohen’s d = 0.20, 0.50, and 0.80 is used to indicate the effect sizes as small, medium, or large, respectively).

Reactive astrocytes increase secretion of key ECM scaffold proteins such as fibronectin. This glycoprotein can also interact with other ECM proteins (i.e., collagen I and laminin) further promoting matrix crosslinking and, thus, contribute to the stiffening and fibrosis of the ONH tissue [15, 35, 36]. As such, we investigated how compressive strain impacts astrocyte secretion of fibronectin. Across all groups, strain increased fibronectin deposition in MONHA-encapsulated hydrogels (Fig. 5). After 4h compression, hydrogels showed a ∼2.5-3-fold increase in fibronectin immunolabeling within the core, and a ∼1.5-fold increase within the periphery (Fig. 5 A, B). Similar trends were observed with 24h compression, such that the core displayed a ∼6-fold increase (Fig. 5 A, C), and the periphery of the ECM hydrogels had a less robust ∼2-fold increase in fibronectin immunolabelling (Suppl. Fig. 8).

**Fig. 5.**
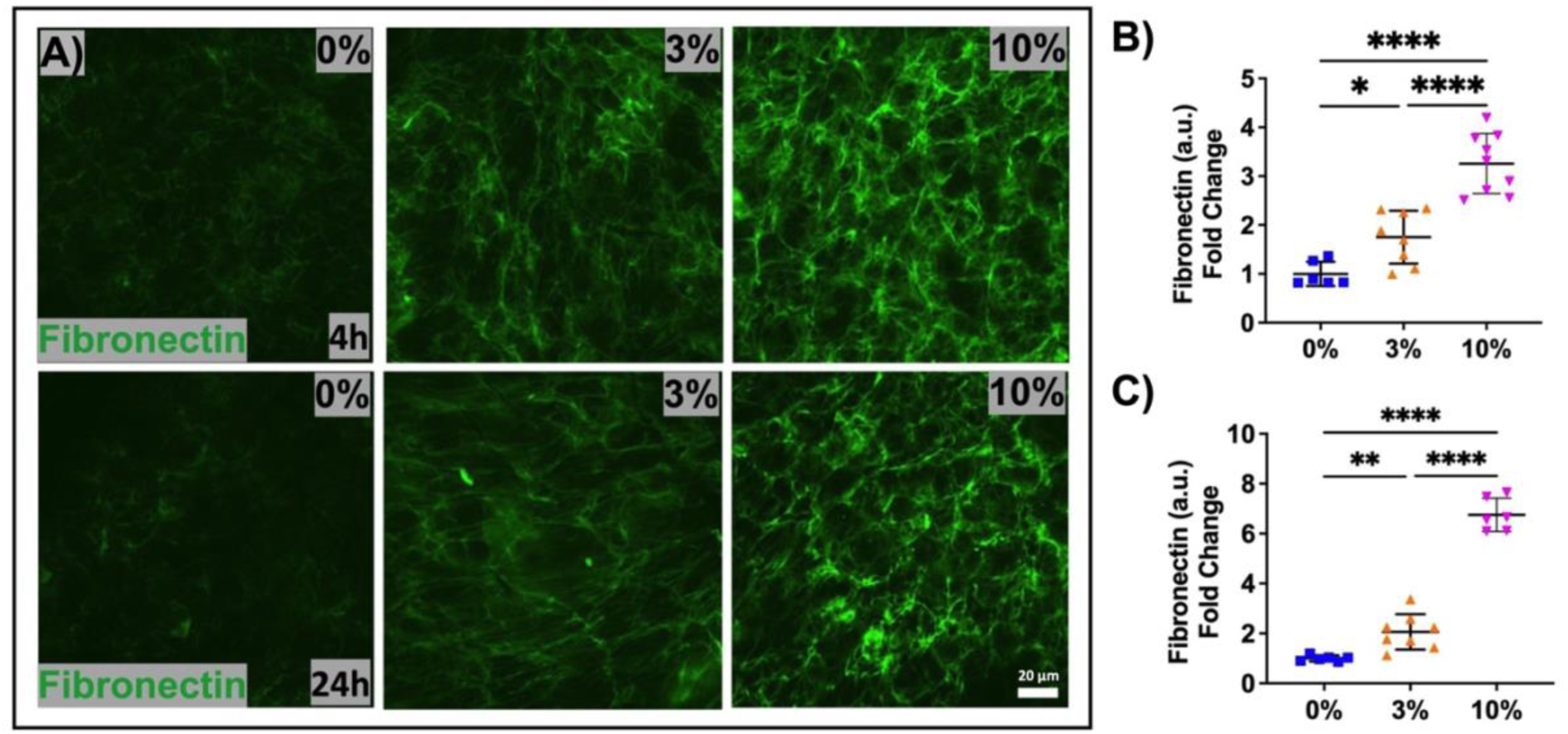
Fibronectin levels in the core of MONHA-encapsulated hydrogels in response to compressive strain. **(A)** Representative fluorescence images of fibronectin in response to duration (i.e., 4h or 24h) and magnitude of strain (i.e., 3% or 10%) (n = 6-9 fields of view/group collected from N = 3 hydrogels/group) and **(B-C)** quantification of fold change in fibronectin signal intensity. Statistical significance was determined using one-way ANOVA (*p < 0.05, **p < 0.01, **** p < 0.0001) for fibronectin signal intensity. Scale bar: 20 μm.

**Supplemental Fig. 8.**
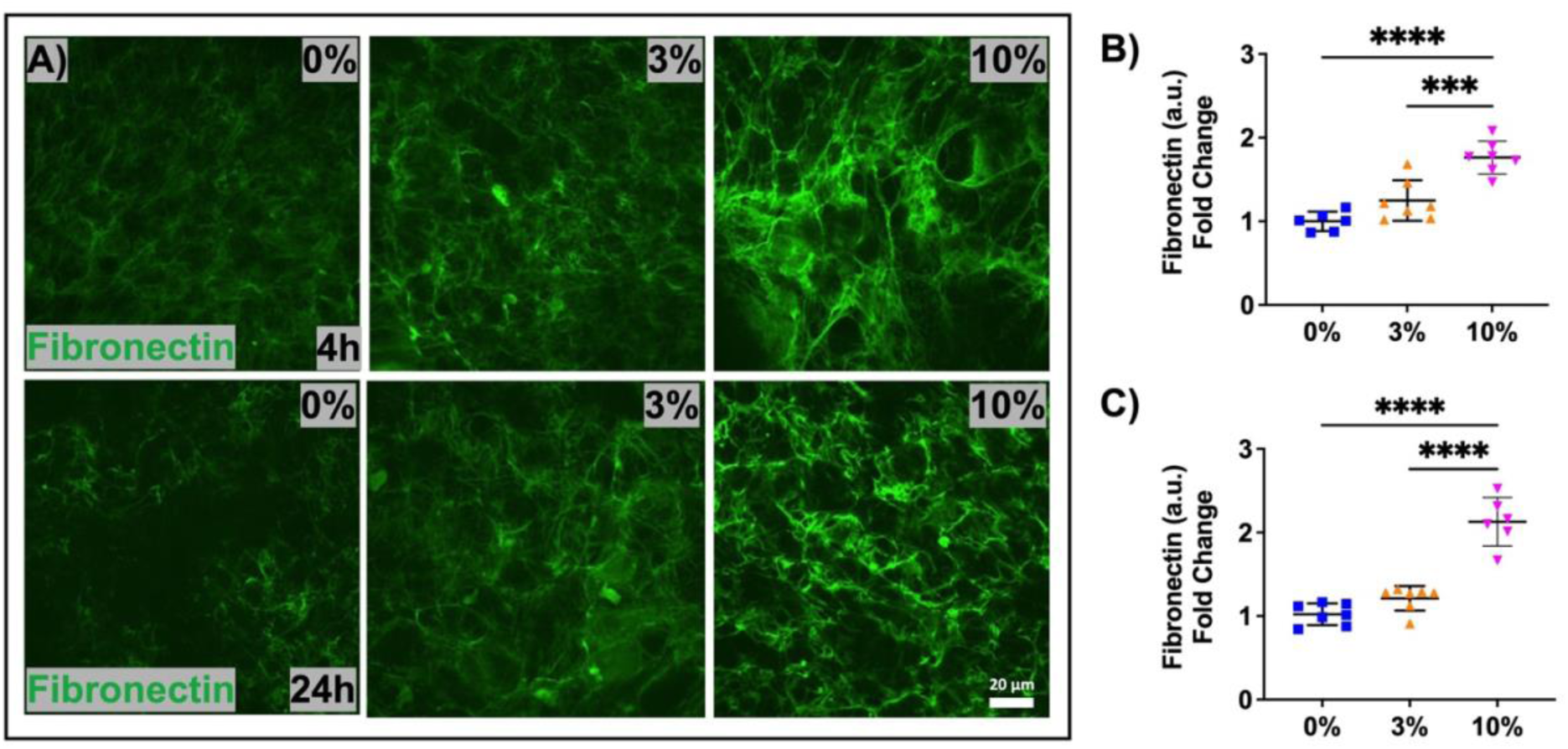
Fibronectin levels in the periphery of MONHA-encapsulated hydrogels in response to compressive strain. **(A)** Representative fluorescence images of fibronectin in response to duration (i.e., 4h or 24h) and magnitude of strain (i.e., 3% or 10%) (n = 6-7 fields of view/group collected from N = 3 hydrogels/group) and **(B-C)** quantification of fold change in fibronectin signal intensity. Statistical significance was determined using one-way ANOVA (***p < 0.001, **** p < 0.0001) for fibronectin signal intensity. Scale bar: 20 μm.

The observed alterations in ECM protein compaction and deposition could potentially impact the global stiffness of MONHA-encapsulated hydrogels. Thus, we analyzed the overall rheological properties of hydrogels after compressive strain. After application of 3% or 10% compression for 24h, we found no significant differences in hydrogel stiffness compared to their uncompressed (0%) controls (Suppl. Fig. 9).

**Supplemental Fig. 9.**
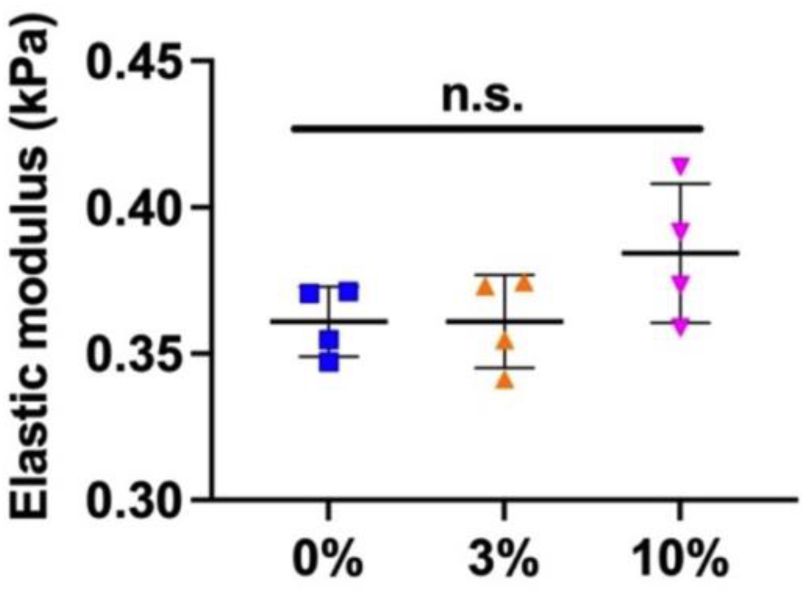
MONHA-encapsulated hydrogel stiffness following compressive strain. Analysis of the elastic modulus (kPa) of MONHA-encapsulated hydrogels (N = 4 hydrogels/group) following 24h compression (i.e., 3% or 10%). Significance was determined by one-way ANOVA using multiple comparisons test.

Collectively, these data show that compressive strain significantly alters collagen distribution and astrocyte fibronectin production in MONHA-encapsulated hydrogels, indicating a dynamic interplay between mechanical stimuli, ECM remodeling, and astrocyte mechanoresponse.

### 2.6 Compressive strain regulates global transcriptomic response of MONHAs across strain level and time

Astrocytes within the ONH are transcriptionally responsive to alterations in IOP. Several murine experimental glaucoma studies have investigated the effects of elevated IOP on ONH astrocyte gene expression and downstream responses [37, 38]. However, it is not clear how glaucoma-associated *biomechanical strains* directly affect ONH astrocyte gene expression and molecular pathways. Given the robust phenotypic response to compressive strain, we next investigated the transcriptional response of ONH astrocytes to compression across strain intensities and durations. Overall, all the RNA samples showed high quality indicated by an RNA integrity number (RIN) ≥ 8.

To assess the global transcriptional effects of compression on astrocytes, we performed principal component analysis (PCA) on bulk RNA sequencing data from MONHA-encapsulated hydrogels subjected to 0%, 3%, or 10% compression for either 4 hours or 24 hours per ECM hydrogel region (i.e., core or periphery). PCA revealed that the duration of compression primarily drove transcriptional variation, with samples separating along PC1 (Suppl. Fig. 10A, B). Strain dynamics in both regions were most observed with 10% strain revealing an initial significant shift along PC2 and PC3 in the 4h groups and maintained in 24h groups.

**Supplemental Fig. 10.**
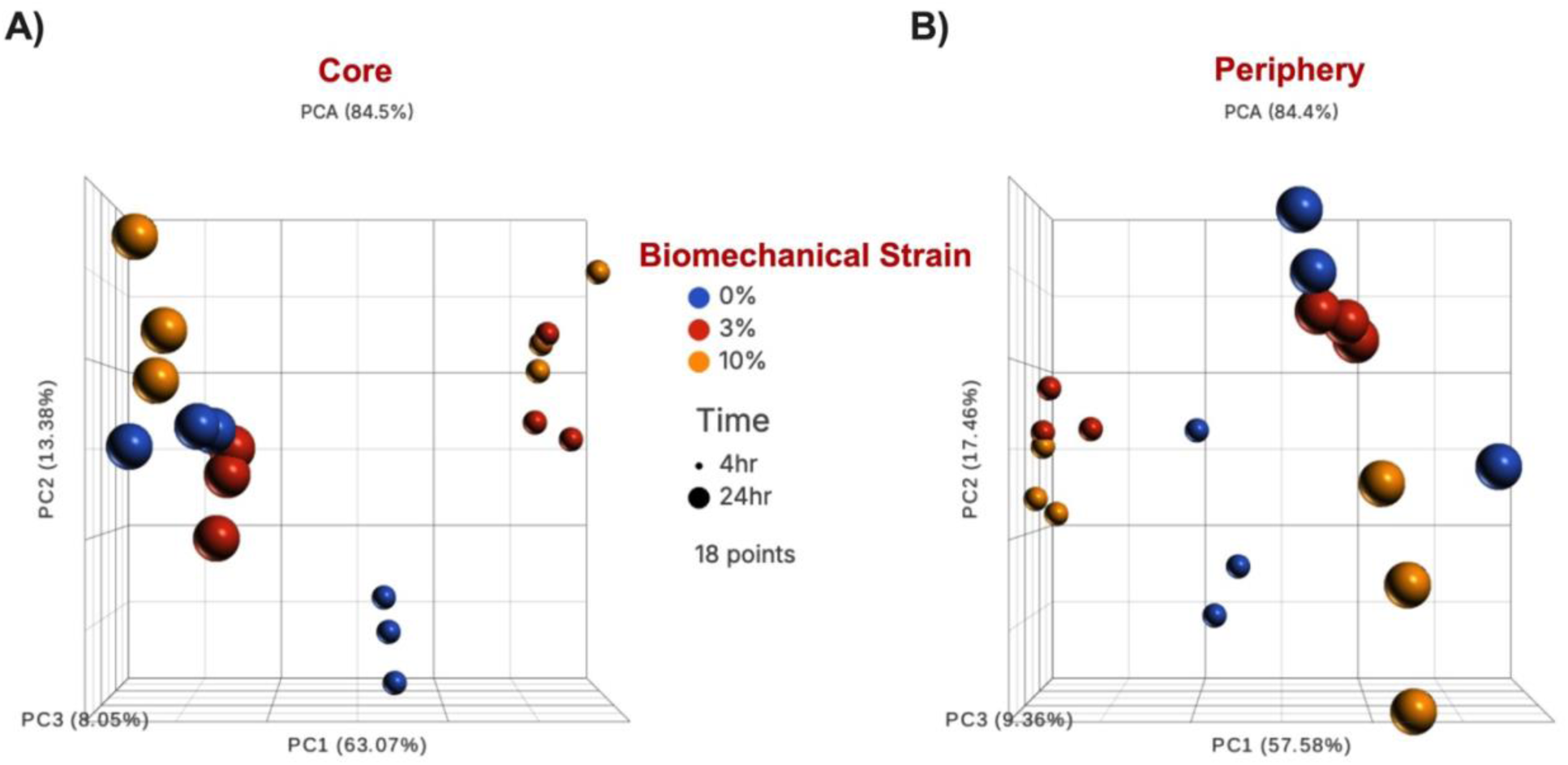
Regional principal component analyses of encapsulated MONHAs transcriptomes in response to compressive strain over time. Each point represents an individual replicate per region (n = 18/core and 18/periphery) subjected to compression (blue = 0%, red = 3%, orange = 10%) for 4 hours (small spheres) or 24 hours (large spheres). Duration of strain (i.e., time) is the dominant source of variation, with point separation along PC1, and magnitudes of strain are observed with PC2 and PC3. **(A)** In the core region, the three principal components (i.e., PC1: 63.07%, PC2: 13.38%, PC3: 8.05%) collectively explain 84.5% of the total variance. **(B)** In the periphery, the three principal components (i.e., PC1: 57.58%, PC2: 17.46%, PC3: 9.36%) account for 84.4% of total variance.

### 2.7 Pairwise analysis reveals distinct transcriptional responses dependent on magnitude of compressive strain and hydrogel region

We next analyzed how different compressive strain intensities (i.e., 3% or 10%) affect MONHA gene expression at different timepoints (i.e., 4h or 24h). DESeq2 analysis identified 19,163 genes after 4h compression and 18,376 genes after 24h compression. We applied a cutoff of FDR ≤ 0.05 and fold change |FC| ≥ 1.5. DEGs from the core of the hydrogel are depicted in Table 1 and in Fig. 6 A, B and Fig. 7 A, B. To simplify the discussion, we have annotated groups by “magnitude of strain” followed by “hydrogel region”; thus, the ECM hydrogel core that underwent 3% biomechanical strain is denoted as *3c*, while the hydrogel periphery is denoted as *3p*. In response to both intensities of biomechanical strain, we observed fewer DEGs in the 24h group vs the 4h group, supporting that encapsulated MONHAs adapt to compressive strain with increasing durations (Table 1).

**Fig. 6.**
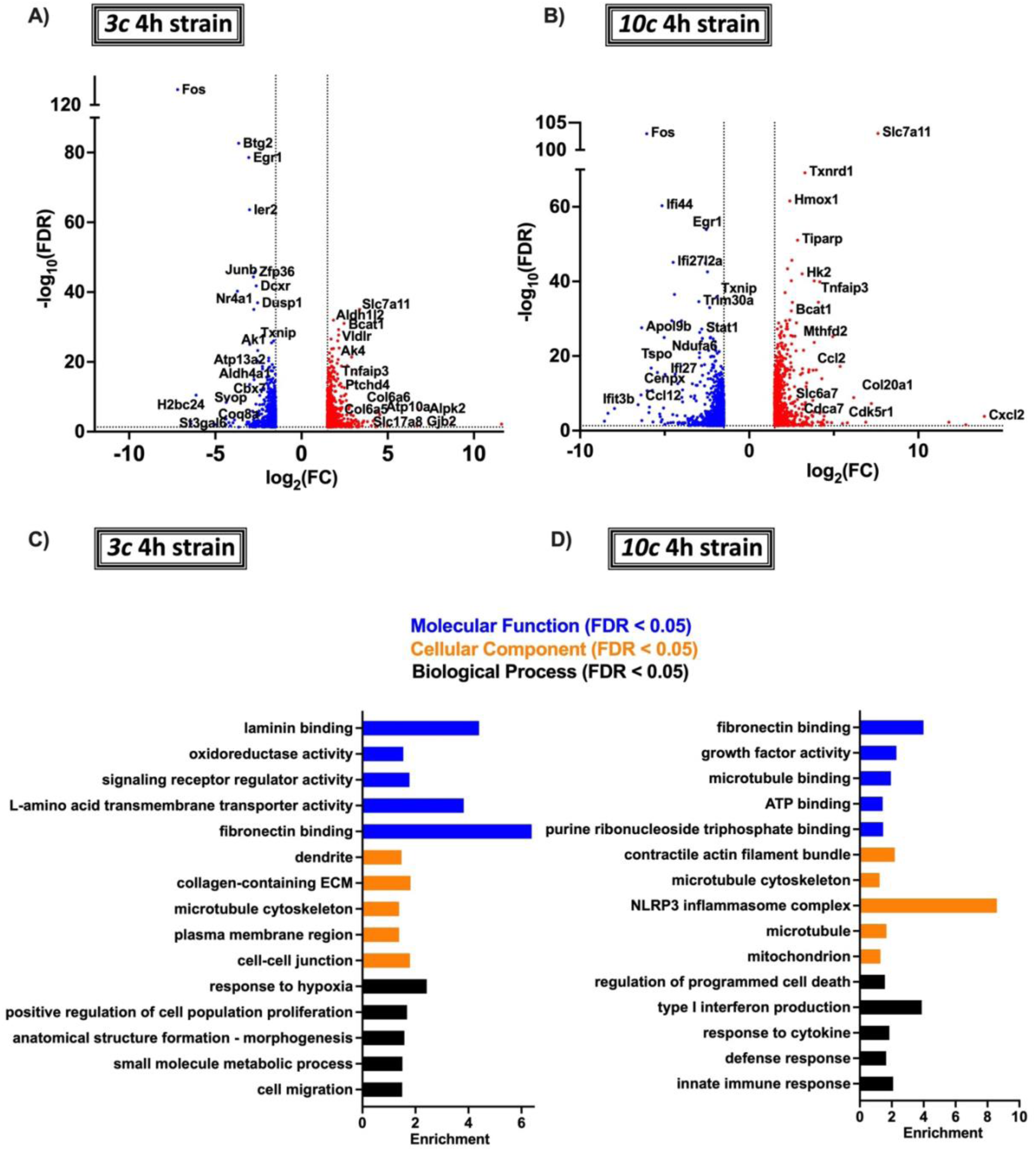
Pairwise transcriptional mechanoresponse of MONHAs encapsulated within the hydrogel core to 4h of compressive strain. **(A-B)** Volcano plots representing the number of DEGs with FDR ≤ 0.05 and |FC| ≥ 1.5 in response to 3% and 10% compression for 4h. **(C-D)** Listed are five representative gene ontology (GO) pathways per functional group of the top 30 most significant GOs at each timepoint.

**Fig. 7.**
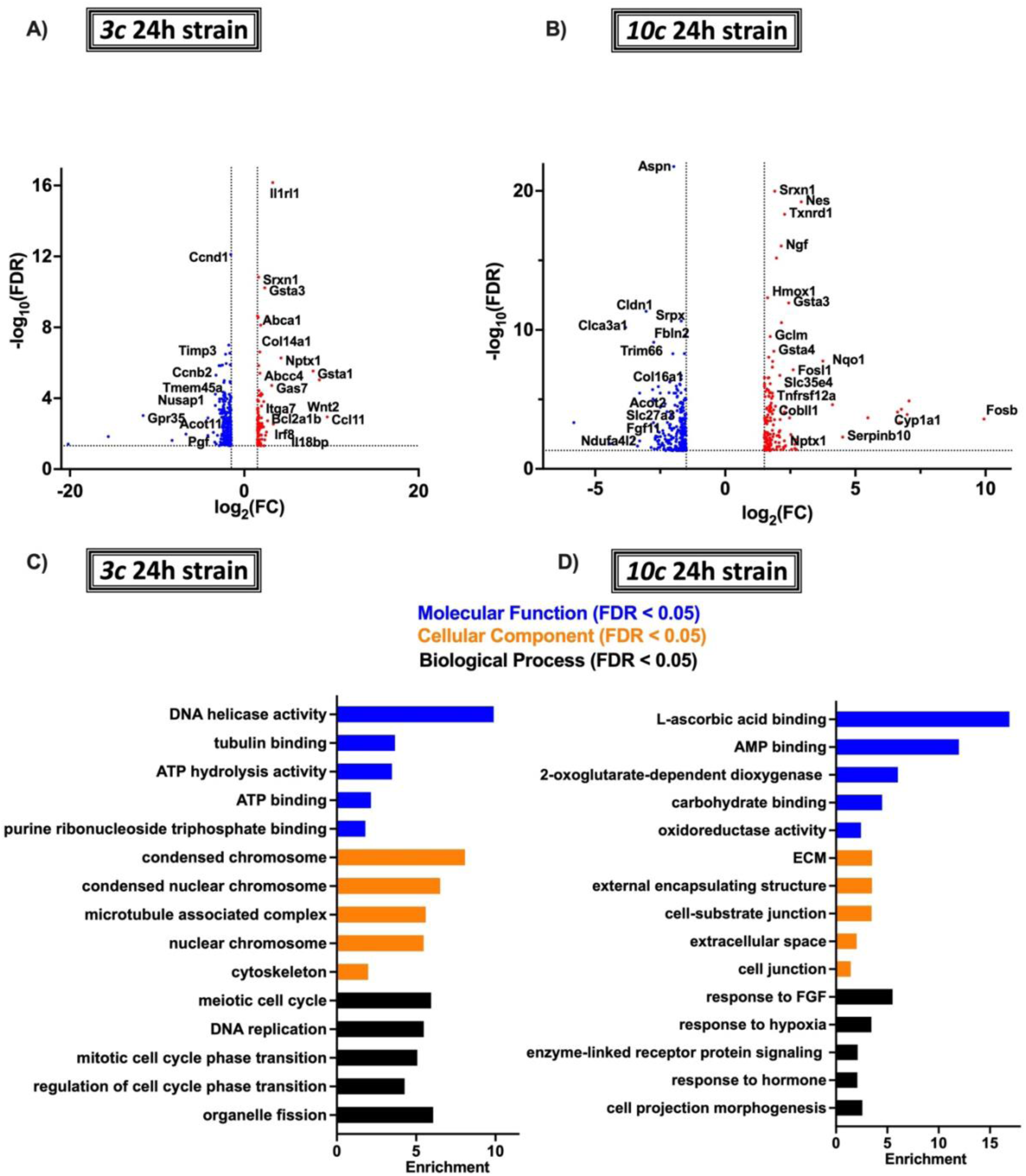
Pairwise transcriptional mechanoresponse of MONHAs encapsulated within the hydrogel core to 24h of compressive strain. **(A-B)** Volcano plots representing the number of DEGs with FDR ≤ 0.05 and |FC| ≥ 1.5 in response to 3% and 10% biomechanical strain for 24h. **(C-D)** Listed are five representative gene ontology (GO) pathways per functional group of the top 30 most significant GOs at each timepoint.

**Table 1.** DEGs with FDR ≤ 0.05 and fold change |FC| ≥ 1.5, core of the hydrogel.

|  | 4h | 24h |
| --- | --- | --- |
| 3% vs 0% | **1,368 total DEGs**<br>686 upregulated, 682 downregulated | **377 total DEGs**<br>104 upregulated; 273 downregulated |
| 10% vs 0% | **1,968 total DEGs**<br>880 upregulated; 1088 downregulated | **413 total DEGs**<br>147 upregulated; 266 downregulated |

For all groups, to identify the functional categories the DEGs were enriched in, the list of DEGs per comparison were uploaded to the publicly available WEB-based GEne SeT AnaLysis platform. In the *3c* group after 4h compression, over representation analysis (ORA) revealed enrichment of GOs involved in cell-cell junction, collagen-containing ECM, laminin/fibronectin binding, and cell proliferation and migration (Fig. 6C). With increased magnitude of strain, the *10c* group after 4h compression showed enriched pathways in the regulation of immune/defense response, response to cytokine, mitochondrion, adenosine triphosphate (ATP)/nucleotide binding, and regulation of programmed cell death (Fig. 6D). These findings suggest that, for relatively short durations (i.e. 4h), increasing the magnitude of compressive strain induced a shift in DEGs enriched in morphology, ECM regulation, and cell proliferation (3%) to enrichment in inflammation regulating and cell cycle/division-related pathways (10%) (Fig. 6C, D).

Similar analyses were done for MONHAs undergoing compressive strain for 24h. For the *3c* group after 24h compression, GOs showed genes enriched in cell cycle, DNA replication, organelle fission, condensed chromosome, microtubule associated complex, DNA helicase activity and ATP binding (Fig. 7C). In contrast, in the *10c* group after 24h compression, GOs showed enriched genes in pathways of carbohydrate/AMP binding, response to hormone, cell–substrate junction, response to hypoxia, and oxidoreductase activity (Fig. 7D). These findings suggest that application of compressive strain for longer durations (24h) induces a shift in DEGs enriched cell cycle and division pathways (3%) to enrichment in metabolism and ECM regulatory pathways (10%) (Fig. 7C, D).

We then analyzed the periphery of the hydrogel, where cells were surrounded by more densely packed ECM (Suppl. Fig. 2). Again, we applied a cutoff of FDR ≤ 0.05 and fold change |FC| ≥ 1.5; DEGs from the periphery of the hydrogel are depicted in Table 2 and in Suppl. Fig. 11 A, B and Suppl. Fig. 12 A, B.

**Table 2.** DEGs with FDR ≤ 0.05 and fold change |FC| ≥ 1.5, periphery of the hydrogel.

|  | 4h | 24h |
| --- | --- | --- |
| 3% vs 0% | **688 total DEGs**<br>336 upregulated, 352 downregulated | **13 total DEGs**<br>6 upregulated; 7 downregulated |
| 10% vs 0% | **801 total DEGs**<br>322 upregulated; 479 downregulated | **445 total DEGs**<br>112 upregulated; 333 downregulated |

In this peripheral region, for relatively short duration compression (4h), we similarly observed that MONHAs displayed a shift in DEGs enriched in cellular signaling regulation (*3p*) to inflammation and metabolism pathways (*10p*) (Suppl. Fig. 11C, D). Interestingly, analysis of the *3p* group after 24h compression identified few DEGs with no significant GOs by our FDR cutoff values. The *10p* group after 24h compression showed GOs primarily enriched in ECM regulation and adhesion (Suppl. Fig. 12C). Taken together, these transcriptomic analyses reveal that astrocytes exhibit highly dynamic time– and region–dependent transcriptional responses to compressive strain, transitioning from early changes in signaling, inflammation and metabolism to later changes in matrix remodeling and adhesion.

**Supplemental Fig. 11.**
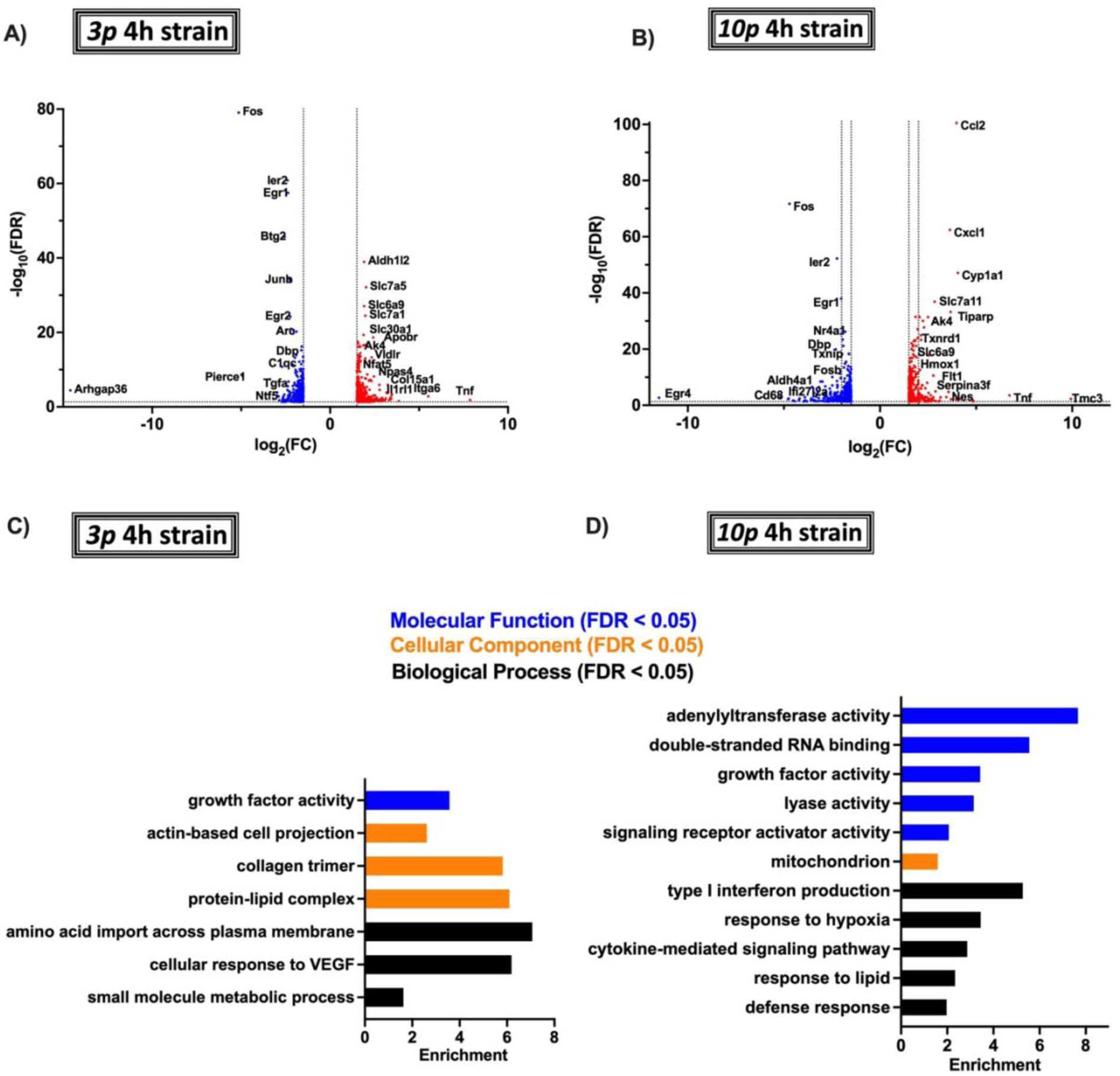
Pairwise transcriptional mechanoresponse of MONHAs encapsulated within the hydrogel periphery to 4h of compressive strain. **(A-B)** Volcano plots representing the number of DEGs with FDR ≤ 0.05 and |FC| ≥ 1.5 in response to 3% and 10% biomechanical strain for 24h. **(C)** Listed are up to five representative gene ontology (GO) pathways per functional group of the top 30 most significant GOs after 4h of strain.

**Supplemental Fig. 12.**
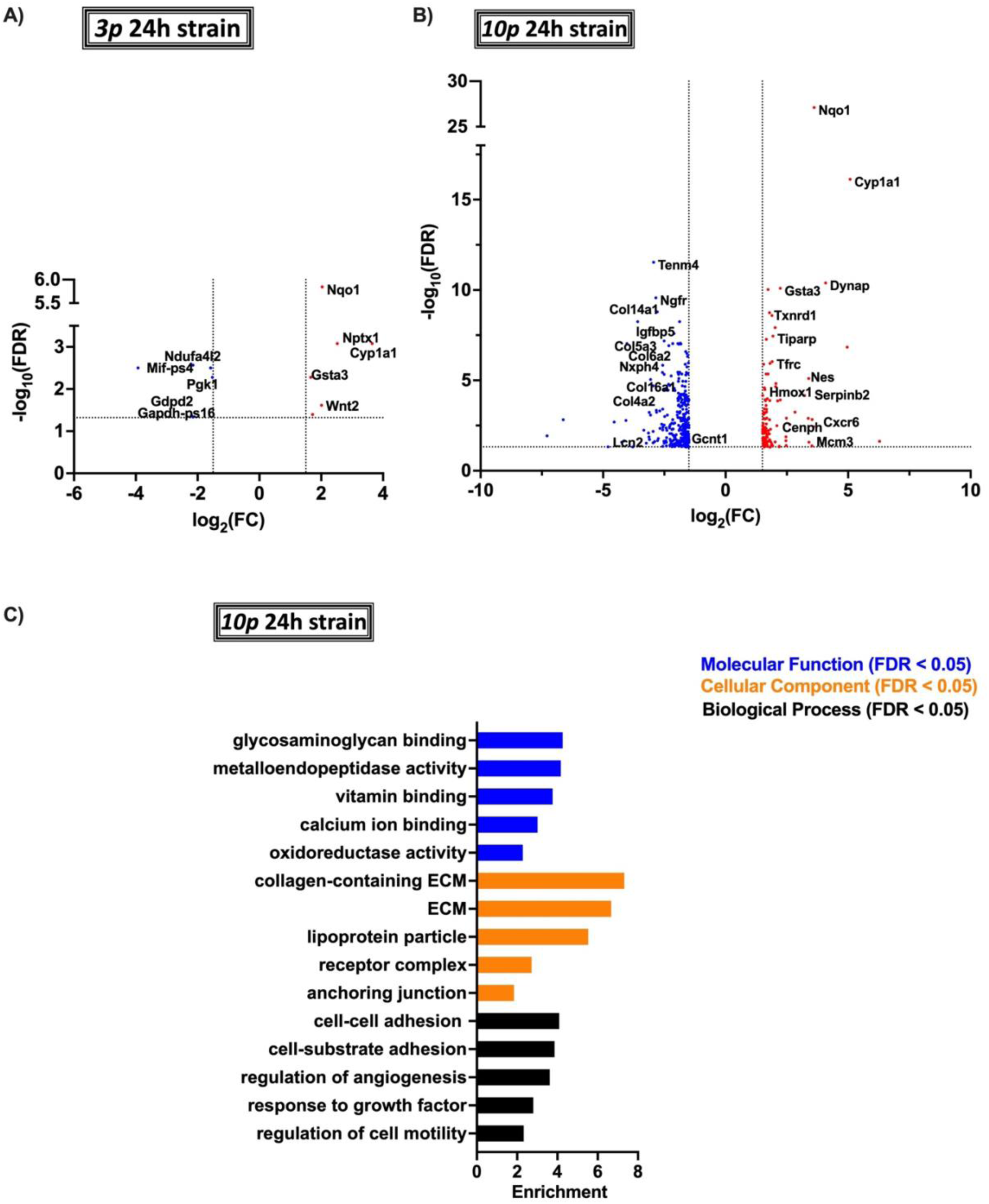
Pairwise transcriptional mechanoresponse of MONHAs encapsulated within the hydrogel periphery to 24h of compressive strain. **(A-B)** Volcano plots representing the number of DEGs for 24h with FDR ≤ 0.05 and |FC| ≥ 1.5 in response to 3% and 10% biomechanical strain for 24h. **(C)** Listed are five representative gene ontology (GO) pathways per functional group of the top 30 most significant GOs at each timepoint.

### 2.8 Hydrogel region and duration of compressive strain modulate key astrocytic molecular and metabolic pathways

To investigate the contribution of increasing magnitude and duration of compression to MONHA transcriptional response, we performed ANOVA of compressive strain groups across timepoints followed by GSEA against both KEGG and Reactome databases. In the <u>hydrogel core</u>, encapsulated astrocytes downregulated pathways related to focal adhesions (*Col6a6; Flnc; Flt1; Hgf; Itga10*), integrin cell surface interactions (*Itga5; Itga7; Itgb3; Itgb8; Lama2*), and ECM interactions (*Col6a5; Col6a6; Itga10; Tnxb*) (Fig. 8A, B, D). These altered pathways corroborate our findings on genes enriched in GOs associated with cell morphology and ECM reorganization. Compressive strain on ECM hydrogel-encapsulated astrocytes also led to the downregulation of key signaling pathways regulating astrocyte quiescence, including those regulating stem cell pluripotency (*Rest; Tbx3; Wnt11*), calcium signaling (*Camk2a; Fzd3; Fzd4*), interleukin-6 (IL6) (*Clcf1; Cntfr; Il6ra; Lif; Lifr; Osmr*), phosphoinositide 3-kinase Ak strain transforming (PI3K-Akt) (*Bcl2l11; Chuk; Vegfa; Flt3l; Gdnf; Hgf; Il6ra*), and Wnt signaling (*Wnt11; Wnt5a*) (Fig. 8A, B, D).

**Fig. 8.**
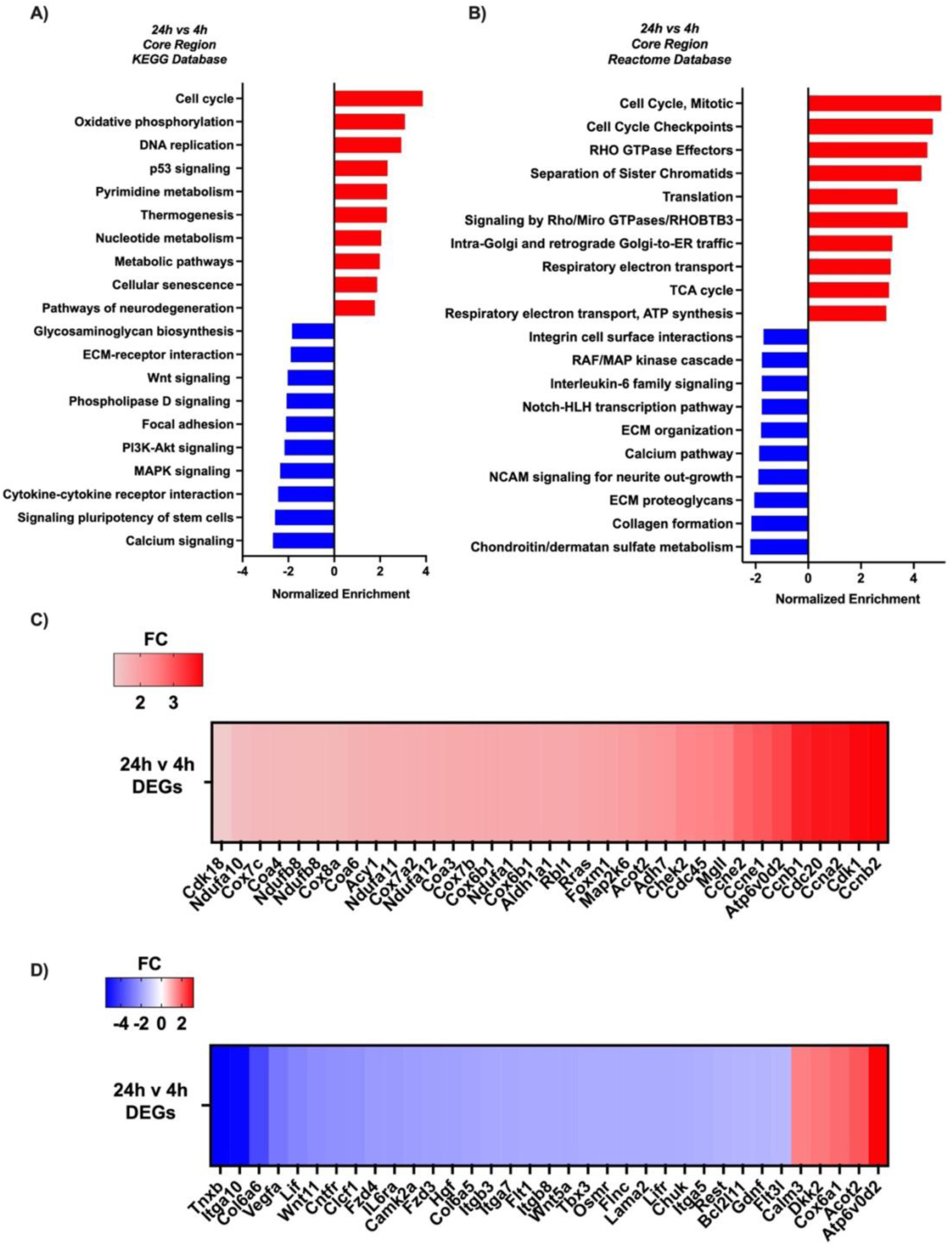
Transcriptional mechanoresponse of MONHAs encapsulated within the hydrogel core over time with increasing compressive strain intensity and duration. **(A-B)** Listed here are 20 up and down most significant pathways against KEGG and Reactome databases with representative 35 DEGs for the **(C)** upregulated and **(D)** downregulated pathways (FDR ≤ 0.05 and |FC| ≥ 1.5) based on ANOVA analysis of MONHAs mechanoresponse over time (i.e., 24h vs 4h) with increasing strain.

**Supplemental Fig. 13.**
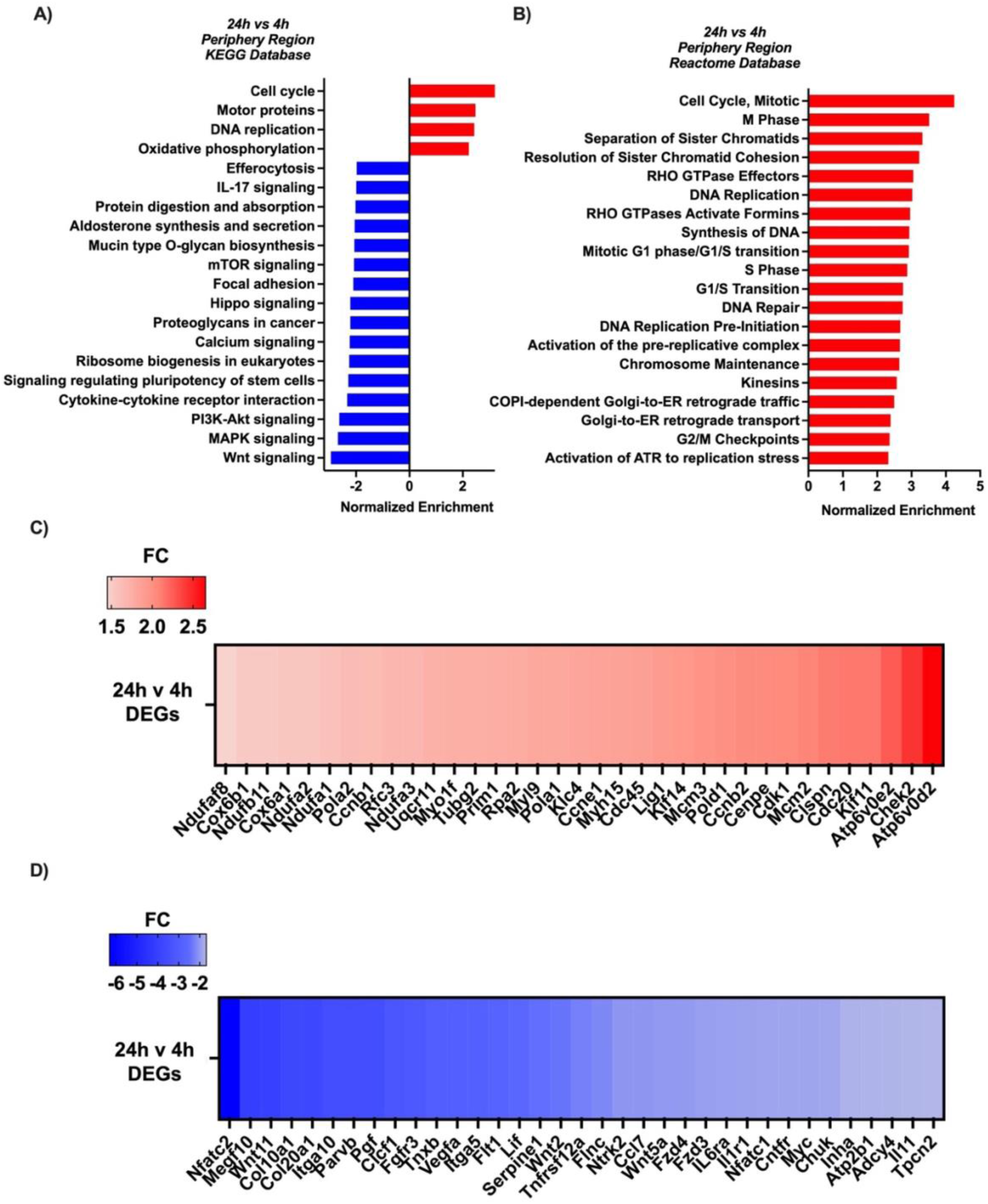
Transcriptional mechanoresponse of MONHAs encapsulated within the hydrogel periphery over time with increasing compressive strain intensity and duration. **(A-B)** Listed here are 20 up and down most significant pathways against KEGG and Reactome databases with representative 35 DEGs for the **(C)** upregulated and **(D)** downregulated pathways (FDR ≤ 0.05 and |FC| ≥ 1.5) based on ANOVA analysis of MONHAs mechanoresponse over time (i.e., 24h vs 4h) with increasing strain.

Additionally, MONHAs encapsulated within the core of the ECM hydrogel showed significant changes in metabolism with increasing time and magnitude of compression. Specifically, we observed upregulated pathways related to metabolic and respiratory electron transport chain (ETC) (Fig. 8A, B, C). As a part of the mitochondrial oxidative phosphorylation, the ETC is crucial for ATP production. These upregulations corresponded with gene enrichment in metabolism/ETC complexes and in thermogenesis (*Acot2; Aldh1a1; Acy1; Adh7; Atp6v0d2; Cox6b1; Cox7a2; Cox7b; Cox7c; Cox8a; Ndufa1; Ndufa11; Ndufa12; Ndufb8*) (Fig. 6A-C). Interestingly, pathways for both cell cycle/repair (*Ccnb1; Ccnb2; Ccne1; Cdc20; Cdc45; Cdk1; Cdk18; Chek2*) and cellular senescence (*Ccna2; Ccne2; Foxm1; Map2k6; Rbl1; Rras*) were also upregulated (Fig 8A-C), potentially indicating a cytoprotective stress response to increasing biomechanical strain over time.

We then investigated the <u>hydrogel periphery</u>, where astrocytes were surrounded by a denser ECM at baseline. Interestingly, we observed a somewhat blunted transcriptional response to compressive strain in the periphery with fewer DEGs (Table 2) identified compared to the core of the hydrogel (Table 1). Similarly to the core region, peripheral MONHAs downregulated pathways related to inflammation such as IL17–signaling, cytokine–cytokine receptor interaction (*Lif; Cntfr; Il11; Il1r1*) as well as signaling pathways implicated in astrocyte quiescence like stem cell pluripotency, calcium or PI3K-Akt/Wnt signaling (*Nfatc2; Tnxb; Vegfa; Wnt2; Wnt5a*) (Suppl. Fig. 13A-D). Importantly, unlike the core, we observed a larger number of significant molecular pathways upregulated in cell cycle, mitosis, DNA synthesis, replication, and repair, ATP activation in response to replication stress, and chromosome maintenance (*Ccnb2; Cdc20; Atp6v0d2; Cox6b1; Ndufa1; Cox6b1; Mcm3*) (Suppl. Fig. 13A-D). As such, the astrocytes within the ECM hydrogel periphery exhibit a distinct shift toward pathways supporting proliferation, DNA maintenance, and replication stress responses. Collectively, these findings show that compressive strain plays a crucial role in shaping astrocyte morphology, function, and ECM regulation concomitant with a robust mechanotranscriptional response.

## 3. Discussion

In glaucoma, increased biomechanical strains on the ONH have long been associated with elevated IOP and disease progression [18–20, 30]; and yet, the mechanoresponse of ONH resident cells remains poorly understood. ONH astrocytes have emerged as key regulators of retinal ganglion cell axonal health. In *in vivo* glaucoma models, these local astrocytes are amongst the first to respond by reorganizing their cytoskeleton architecture and gap junction coupling [12–14, 27, 39]. In the present study, we sought to comprehensively analyze the direct phenotypic and transcriptional responses of astrocytes to glaucomatous biomechanical strains, to better understand the role of these pathologic biomechanical forces on astrocyte behavior.

### Dissecting the ONH astrocyte mechanoresponse by modeling glaucomatous compressive strains in a 3D ECM hydrogel

Prior studies explored ONH astrocyte mechanoresponse to glaucomatous biomechanical strains using conventional 2D *in vitro* cultures [17, 40]. However, it is increasingly recognized that the ECM significantly influences cell behavior, with mechanical, structural, and compositional cues playing critical roles [41]. Newer *in vitro* 3D models, including our own [22], utilize viscoelastic ECM biopolymer hydrogels made from collagen and hyaluronic acid to better mimic the 3D architecture and promote a more natural astrocyte morphology [6, 41, 42]. By encapsulating astrocytes within a collagen/hyaluronic acid network, one can directly visualize cell–cell and cell– ECM interactions. We previously showed that hydrogel-encapsulated murine ONH astrocytes extend processes in 3D, couple with astrocyte neighbors, and bind to and modify their surrounding ECM [22]. In this study we sought to comprehensively investigate how 3D-encapsulated ONH astrocytes respond to glaucoma-related compressive strains over time. A custom bioreactor previously used on collagen-encapsulated rat cortical astrocytes [23], was used to apply increasing intensities of compressive strain over time to investigate nuanced ONH astrocyte phenotypic and transcriptomic responses to mechanical stimuli.

### Compressive strain triggers a multifaceted phenotypic and transcriptomic response in encapsulated astrocytes, mimicking aspects of glaucomatous ONH astrocyte dysfunction

Four main astrocyte mechanoresponses were observed in this study. After compression, MONHAs demonstrated alterations in (i) cell division/senescence, (ii) cytoskeletal architecture and morphology, (iii) ECM remodeling, and (iv) metabolic function. These findings are largely in line with ONH astrocyte behavior in *in vivo* glaucoma models, and highlight the potentially instrumental role of glaucomatous compressive strains in regulating ONH dysfunction. Our finding of robust transcriptional alterations related to cell cycle regulation and DNA replication suggests development of a proliferative and/or senescent phenotype in MONHAs following compression. Early in *in vivo* glaucoma models, ONH astrocytes proliferate, and exhibit increased reactive gliosis [11, 39, 43]. With prolonged IOP elevation, ONH astrocytes show evidence of cell senescence and death [44, 45]. Interestingly, in our study, transcriptional alterations did not correlate with altered astrocyte viability in response to relatively short durations of compressive strain (i.e., 4h and 24h), suggesting that longer durations compressive strain are necessary to induce astrocyte senescence/death in 3D *in vitro*.

One of the most profound findings in our study was a robust change in F-actin cytoskeleton architecture after compressive strain. In the core of the ECM hydrogel, there was a significant decrease in F-actin area coverage, suggestive of MONHA process retraction that correlated with increasing intensity and duration of compression. We also observed downregulated pathways of focal adhesion and ECM-interactions, and significant GOs pertaining to cytoskeleton, actin–based cell projection, dendrite and cell–cell junction. This is particularly interesting in light of *in vivo* evidence that ONH astrocytes undergo a dramatic reorientation of their F-actin cytoskeleton within 8 – 72 h of acute IOP elevation [10, 13, 14, 39], and a decrease in astrocyte process complexity and coupling [27, 44]. These early morphologic changes in ONH astrocytes may reduce astrocyte– astrocyte connectivity as well as contact with surrounding blood vessels, retinal ganglion cell axons, and ECM, potentially propagating glaucomatous damage.

Moreover, the observed morphologic alterations corresponded with reorganization of the surrounding ECM. After 24h compression, there was a significant reduction in collagen compaction surrounding MONHAs in the core of the hydrogel, suggesting that the changes in F-actin area coverage may diminish cell–ECM interactions, thereby hindering their ability to remodel surrounding collagen fibrils. We also observed increased fibronectin levels and significant enrichment of genes associated with collagen, fibronectin, and ECM modulation. This response of MONHAs to pathologic compressive strain aligns with the behavior of ONH astrocytes *in vivo*; in glaucoma, astrocytes modify the ECM extensively, leading to increased ECM protein (such as fibronectin) production and crosslinking [15, 30, 35, 36, 46, 47].

A key feature of the glaucomatous ONH is the dysregulation of metabolic pathways [30, 31, 39, 48, 49]. Within 4h compression, there was increased HIF-1α expression, which can trigger transcription of hypoxia–responsive genes, affecting downstream metabolism [34, 50, 51]. Simultaneously, sequencing revealed enrichment of genes involved in cell response to hypoxia pathways, which persisted at longer durations (24h). We also observed upregulated metabolic and respiratory electron transport/ATP synthesis pathways with genes enriched in mitochondrion and ATP binding. In glaucoma models, the optic nerve undergoes hypoxia and metabolic insufficiency, correlating with increased glycolysis and ATP production [34, 52–54]. In response to compressive strain, we observed similar early (within 4h) alteration in pathways related to hypoxia and oxygen levels.

We were surprised to find certain divergent phenotypes in the periphery of the ECM hydrogel when compared to the core. In the periphery, F-actin area coverage and cellular/process morphology remained unchanged after compressive strain, and we observed a surrounding *increase* in collagen compaction. Additionally, while compressive strain resulted in gene enrichment in cell cycle and numerous ECM regulatory pathways (similar to the astrocytes residing in the core), there was greater gene enrichment in metabolic pathways such as mitochondrion, vitamin binding, small metabolic process, oxidoreductase activity, and response to growth factors. Notably, there were numerous pathways upregulated in cell cycle/stress, oxidative phosphorylation and organelle protein trafficking, and downregulated reactivity/inflammation. At baseline (prior to application of compressive strain), this peripheral region was characterized by increased collagen compaction (Suppl. Fig. 2). It is possible that a denser ECM in the hydrogel periphery may dampen the pro-inflammatory astrocyte phenotype, while increasing cell cycle and metabolic alterations. *In vivo*, acute IOP elevation is associated with significant alteration in metabolism-related genes in ONH astrocytes, with a relatively muted inflammatory response [37, 55]. This aligns with our compression injury-induced *in vitro* model. The regional differences in phenotypic and transcriptomic responses by hydrogel-encapsulated astrocytes highlight the important role that ECM integrity plays in regulating astrocyte response to biomechanical strain.

### Limitations and future directions

The presented *in vitro* ECM hydrogel model provides evidence for compressive biomechanical strain injury as a driving force in ONH astrocyte pathophysiology in glaucoma. While astrocyte mechanoresponse was largely consistent throughout the hydrogel, there were a few regional differences observed. Potential reasons for this variability include baseline differences between the core and periphery of the hydrogel, such as the baseline ECM compaction (as evidenced in Suppl. Fig. 2), relative differences in biomechanical strain profile across regions, and variable oxygen concentration within the ECM hydrogel. Finite element modeling of the strain profile experienced throughout the unconfined hydrogel using this compression bioreactor predicted a relatively uniform compressive strain distribution [24], but the contribution of regional variations in tensile and shear strain is unknown. It is also possible that large hydrogels (i.e., 8 x 4 mm constructs) have decreased oxygen concentration in the core as compared to the periphery of the hydrogel. However, we observed prompt tracer diffuse through the hydrogel (Suppl. Fig. 1), suggesting that cells in the core of the ECM hydrogel have adequate access to media nutrients. Additionally, greater gene enrichment regulating metabolic processes were observed in the periphery of the hydrogel compared to the core, where one would expect relatively higher oxygen concentration.

Here, we encapsulated MONHAs in a relatively simplified fibrillar ECM hydrogel, which does not precisely recapitulate the complex composition of a primate lamina cribrosa nor a murine astrocytic lamina [35, 56]. In the future, the hydrogel could be modified by ECM patterning, to generate a more representative lamina cribrosa–like structure. Many of our insights into mechanisms of glaucoma pathophysiology are gained from experimental rodent models [12, 26, 27, 37, 57]. Building on this foundation, we encapsulated *murine* ONH astrocytes to compare and contrast to the *in vivo* findings. Furthermore, by isolating and precisely controlling application of biomechanical strain, we established a platform to directly dissect the specific contribution of compressive strain to astrocyte mechanoresponses. Our model system does allow for the incorporation of other resident cells, including human ONH cell types or microglia, enabling analysis of different cell–cell interactions under biomechanical strain. Compressive strains of 3% and 10% were selected based on finite element modeling predictions of the strains experienced by the glaucomatous ONH [19, 23, 58, 59], and 1Hz was selected to mimic the cyclic ocular pulse amplitude [60]. Future studies can modify superimposed strain and frequency dependent on the species of origin (i.e., murine cells vs. human cells).

### Conclusions

Glaucoma is a complex multifactorial neurodegeneration, for which pathologic biomechanical strain is a key driving factor in ONH pathology. By encapsulating astrocytes from the tissue of interest (i.e., the ONH) within a biologically-relevant ECM hydrogel, we have comprehensively defined the contribution of these compressive strains to ONH astrocyte behavior. As the key mechanosensors of the ONH, astrocytes are critical for initiating complex molecular cascades that can result in both protective and detrimental outcomes, underscoring the importance of studying their nuanced mechanoresponse.

## 4. Methods

### 4.1 Mouse ONH astrocyte isolation and culture

C57BL/6J mice were purchased from the Jackson Laboratory (Bar Harbor, ME) and bred in house according to the institutional guidelines for the humane treatment of animals (IACUC #473) and to the ARVO Statement for the Use of Animals in Ophthalmic and Vision Research. Each harvest used 5-8 mice aged 6-8 weeks. The isolation/culture of primary mouse ONH astrocytes (MONHAs) was performed as previously described [32]. Briefly, optic nerve head tissue was dissected from each eye using a SMZ1270 stereomicroscope at a location proximal to the sclera. The samples then underwent digestion in 0.25% trypsin (Invitrogen, 25200-056, Carlsbad, CA, USA) for 15 minutes at 37 °C. Following digestion, the tissue was resuspended in MONHA growth medium, consisting of Dulbecco’s modified Eagle’s medium, DMEM/F12 (Invitrogen, 11330-032, Grand Island, NE, USA), supplemented with 10% fetal bovine serum (FBS; Atlanta Biologicals, S11550, Atlanta, GA, USA), 1% penicillin/streptomycin (Corning, 30-001-CI, Manassas, VA, USA), 1% Glutamax (Invitrogen, 35050-061, Grand Island, NY, USA), and 25 ng/ml epidermal growth factor (EGF; Sigma, E4127-5X, St. Louis, MO, USA). The digested ONH tissue was then plated onto T75 cell culture flasks coated with 0.2% gelatin and incubated at 37 °C in a humidified atmosphere with 5% CO2. MONHAs migrated from the ONH tissue over a period of 10-14 days before the first passage. All subsequent experiments included MONHAs at passage 3.

### 4.2 Preparation of PDMS molds

A polydimethylsiloxane (PDMS) mixture was used at a 10:1 ratio of elastomer to curing agent as per the manufacturer’s instructions (PDMS; Sylgard 184, Dow Corning, Midland, MI, USA). This mixture was poured into custom 3D printed ABS-M30 filament negative molds (F170; Stratasys, Eden Prairie, MN, USA), measuring either 10 mm diameter x 6 mm depth or 8 mm diameter x 4 mm depth, cured at 60°C, and autoclaved prior to use.

### 4.3 Preparation of MONHA-encapsulated ECM hydrogels

MONHA-encapsulated hydrogels were prepared as previously described [22]. Briefly, MONHAs (2.5 × 10^6^ cells/ml) in media were mixed with 3.1 mg/ml methacrylate-conjugated bovine collagen type I (MA-COL; molecular weight: ∼300 kDa, degree of methacrylation: ∼60– 70%; Advanced BioMatrix, Carlsbad, CA, USA), 1 mg/ml thiol-conjugated hyaluronic acid (SH-HA; Glycosil®; molecular weight: ∼300 kDa, degree of thiolation: ∼20–30%; Advanced BioMatrix), and 0.025% (w/v) riboflavin (Sigma-Aldrich, R7774, Darmstadt, Germany) (photoinitiator) on ice. The chilled MONHA hydrogel precursor solution was pipetted as 250 μl into 8 x 4 mm PDMS molds or 500 μl into 10 x 6 mm PDMS molds. Constructs were then crosslinked by exposure to low intensity blue light (405-500 nm) (OmniCure S1500 UV Spot Curing System; Excelitas Technologies, Mississauga, Ontario, Canada) at 10.3 mW/cm^2^ intensity for 5 minutes. Following hydrogel crosslinking, each MONHA-encapsulated hydrogel was carefully removed from its respective PDMS mold and left in MONHA growth medium [22]. MONHA-hydrogels were cultured for 14 days prior to any experiments, and media was replenished every 2-3 days.

### 4.4 MONHA-encapsulated ECM hydrogel pore size analysis

The collagen network of acellular and MONHA-encapsulated hydrogels was imaged with confocal reflectance microscopy on a Zeiss LSM980 scanning confocal microscope (Carl Zeiss Microscopy, Jena, Germany) using a 40x water immersion objective (LD LCI Plan-Apochromat 40x/1.2 Imm Corr DIC M27, Zeiss). The excitation wavelength was set to 488 nm and backscatter signal was collected via a beam splitter BC80/20. Pore size within the collagen gel was calculated using an algorithm as previously described [61]. In brief, 6-9 fields of view were obtained from the core and periphery of MONHA-encapsulated hydrogels for each magnitude of strain (i.e., 0%, 3%, or 10%) at each timepoint (i.e., 4 hours, 24 hours). The image size was set to 1024 x 1024 pixels in x/y and the z-step interval set to 0.5 μm collecting a total of 50 μm in the z-axis direction. The confocal z-stack images of the collagen gel were input to FIbeR Extraction MATLAB code [62] to extract the Euclidean distance map. The pore size was then extracted using the bubble analysis algorithm in MATLAB [63].

### 4.5 MONHA-encapsulated ECM hydrogel permeability analysis

To determine the permeability of MONHA-encapsulated hydrogels, acellular and MONHA-encapsulated hydrogels were crosslinked on top of 3 μm pore size transwell inserts in 12-well plates (Thermo Fisher, 141080, Kennebunk, Maine, USA). Before initiating the permeabilization assay, gels were equilibrated in DMEM/F12 for 24h. The assay was initiated by adding 200 μl of a 1000 μg/ml FITC-labelled dextran (70 kDa) tracer solution (FITC-DEAE-Dextran; Sigma, 54702-1G) into the top chamber above each hydrogel. In the bottom chamber, 1000 μl Dulbecco’s Phosphate Buffered Saline 1X (DPBS; Invitrogen, 14190-44) was added which served as receiving medium. The tracer molecule leakage into the bottom well was quantified (i.e., after permeating through the hydrogel). A volume of 600 μl from the bottom well was withdrawn at each time point (5, 10, 15, 30, 45 min – 1, 2 h), and each time it was replenished by an equal volume of fresh DPBS. For quantification, a standard curve was generated using graded dilutions of FITC-labelled dextran (1 to 1000 μg/ml). A volume of 200 µL for each supernatant sample and standard were transferred into a 96-well plate and absorbance was measured at 490 nm using a BioTek Gen5 microplate reader (version 3.16; Agilent Technologies, Palo Alto, CA, United States). The concentration of “leaked” FITC-labelled dextran in the collected samples was determined using a standard curve equation and as previously described [64, 65]. Cumulative FITC-labelled dextran at each time point was calculated by using:

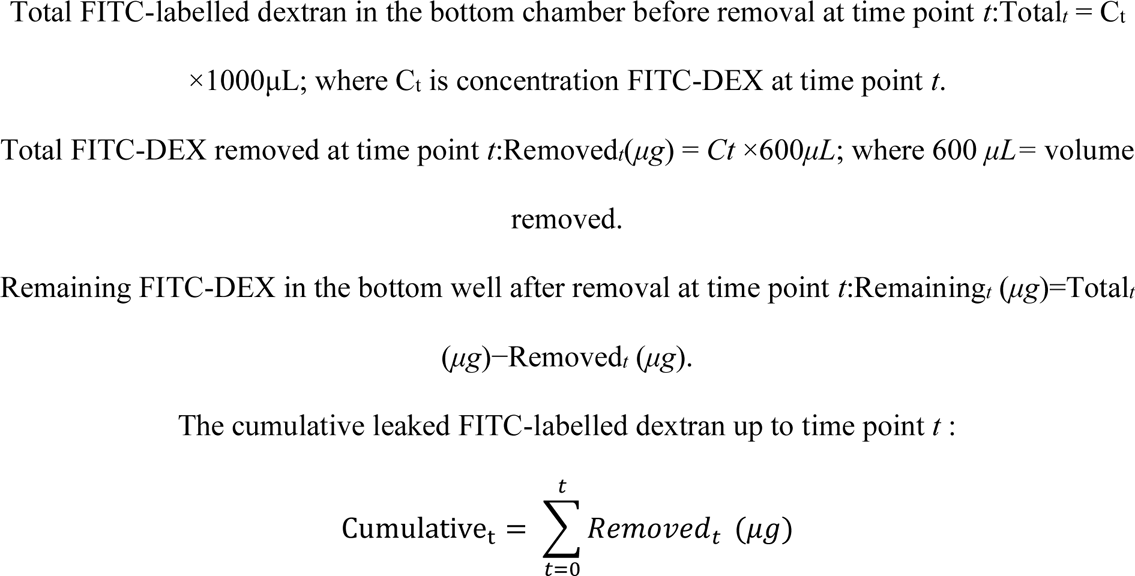

### 4.6 MONHA-encapsulated ECM hydrogel collagen fiber signal intensity, compaction and cell alignment analysis

For collagen signal intensity and compaction analyses, MONHA-encapsulated hydrogels were imaged using confocal reflectance microscopy (see **Methods 4.4**). Fold-change collagen signal intensity in MONHA-encapsulated hydrogels was quantified with background subtraction from an acellular hydrogel using Z-project Maximum Intensity Projection in FIJI and further normalized to number of nuclei. Collagen compaction was determined as previously described [66]. In brief, the mean intensity of collagen fibrils in the fixed area around the astrocytes was computed and then normalized against the mean intensity of the collagen fibrils as deduced from acellular collagen images. The cropped area was found with binarized fluorescent images of the astrocytes with pretrained denoising convolutional neural network (dsCNN)/median filter and adaptive threshold functions in MATLAB. Moreover, due to the large sample size (number of pixel intensity N > 10000), we used Cohen’s effect size d to compare the relative difference between the mean value of different groups (i.e., substantive significance), which scales with sample size unlike p-value (i.e., statistical significance) [67]. Cohen’s d = 0.20, 0.50, and 0.80 was used to indicate the effect size as small, medium, or large, respectively.

### 4.7 Mechanical loading of MONHA-encapsulated ECM hydrogels

Using a custom bioreactor system [23–25], unconfined MONHA-encapsulated hydrogels were cultured for 14 days, then subjected to either 0%, 3% or 10% cyclic compressive strain at 1Hz for 4 hours and 24 hours. These biomechanical strain intensities have been previously correlated with strains within the glaucomatous ONH [18–20]. Hydrogels were placed on PDMS-coated glass containers and maintained in 40 ml of astrocyte media for the duration of biomechanical strain. A thermoplastic polyester platen attached to a linear actuator (Zaber Inc., NA08A30-T4), with 5 mm clearance from the glass wall and 10 perforations (5 mm in diameter/perforation) cyclically compressed the hydrogels to 3 or 10% of their height. 0% static controls were maintained in similar conditions with the platen immediately above the hydrogels, but not actively compressing them. For post-strain analysis, the top 1 mm (adjacent to the compressive platen) and bottom 1 mm (adjacent to the glass container) were discarded as to avoid any biased effects caused by contact with the platen or glass container, and the 3 mm core was further separated from the periphery of the hydrogel to allow for systematic comparisons between different regions, magnitudes and durations of biomechanical strain.

### 4.8 Encapsulated MONHA cell viability analyses

Cell viability within the hydrogel was determined using a CellTiter 96® Aqueous Non-Radioactive Cell Proliferation Assay (MTS; Promega, Madison, WI, USA) as per the manufacturer’s instructions. Following 24 hours of cyclic strain, MONHA-encapsulated hydrogels were incubated with the staining solution (114 μl MTS, 6 μl PMS solution, 600 μl media) at 37 °C for 4h, and absorbance at 490 nm was assessed with a spectrophotometer plate reader (BioTEK, Winooski, VT, USA). Fold-change over time of blank-corrected absorbance values was analyzed to quantify cell viability in hydrogels.

### 4.9 Encapsulated MONHA morphological analysis

Following application of compressive strain, MONHA-encapsulated hydrogels were fixed with 4% PFA at 4 °C overnight, permeabilized with 0.5% Triton^TM^ X-100 (Thermo Fisher Scientific) and stained for filamentous actin (F-actin; Alexa Fluor® 488- or 594-conjugated Phalloidin (1:200, Abcam, Ab176757 or Ab176753); nuclei were counterstained with DAPI. Fluorescent z-stack images were captured using an air 20x objective (Plan-Apochromat 20x/0.8 M27, Zeiss) and a Zeiss LSM780 scanning confocal microscope (Carl Zeiss Microscopy, Jena, Germany). The image size was set to 1024 x 1024 pixels or 512 x 512 in x/y, and six titled z-stack images consisting of 100 μm in z-axis direction with the z-step interval set to 1.3 μm were obtained. Assessment of cell morphology was conducted using Z-project Maximum Intensity Projection in FIJI (NIH, Bethesda, MD) across individual z-stacks. F-actin masks were then generated and binarized in FIJI, and the area of F-actin coverage was calculated by the ratio of percentage of pixels with non-zero pixels [68].

### 4.10 MONHA-encapsulated ECM hydrogel sectioning and immunohistochemical analysis

MONHA-encapsulated hydrogels were cultured for 14 days prior to compressive loading for 4h and 24h and then fixed with 1% Pagano fixative solution (i.e., 4% PFA, 50 mM Sucrose, 25 mM MgCl2, 2.5 mM KCl, HEPES 7.4) [69] at 4°C overnight. Samples were then washed with DPBS and embedded in 10% bovine gelatin (Sigma-Aldrich, G9382-500G). Subsequently, gelatin-embedded MONHA-encapsulated hydrogels were fixed again with 1% Pagano fixative solution at 4°C overnight and then washed with DBPS. Using the Leica VT1000 S vibratome, 150 μm sections were cut and collected in cold PBS. Sections were then permeabilized with 0.5% Triton^TM^ X-100 for 15 min, blocked in 7% normal goat serum (Abcam, Ab7481) for 1h and immunostained in 1% normal goat serum (Abcam, Ab7481) for either GFAP (Anti-GFAP Antibody Cy3 Conjugate, 1:100, Millipore, MAB3402C3), fibronectin (rabbit anti-fibronectin antibody, 1:100, Abcam, Ab2413) or hypoxia-inducible factor 1α (HIF-1α; rabbit anti-HIF-1 alpha antibody, 1:200, Abcam, Ab228649) overnight, followed by stain with Alexa Fluor® 488-conjugated secondary antibody (goat polyclonal antibody to rabbit IgG, 1:500, Abcam, Ab150077) for 1 h; nuclei were counterstained with DAPI (Abcam, Ab228549). Sections were mounted with ProLong^TM^ Gold Antifade (Thermo Fisher Scientific, P36930, Waltham, MA, USA) on Superfrost^TM^ Plus microscope slides (Fisher Scientific, 1255015). 3-4 fields of view per core or periphery of the hydrogel were acquired using an air 40x objective (S Plan Fluor 40x/0.6, Nikon) and an Andor BC43 SR spinning-disk confocal microscope (Oxford Instruments, Belfast, Northern Ireland). The image size was set to 2040 x 2040 pixels in x/y and the z-step interval set to 1 μm collecting a total of 40 μm in the z-axis direction. Fold-changes in normalized signal intensity were quantified with background subtraction using Z-project Maximum Intensity Projection in FIJI and further normalized to number of nuclei for GFAP and HIF-1α analyses.

### 4.11 MONHA-encapsulated ECM hydrogel rheology analysis

MONHA-encapsulated hydrogels, fabricated using the 10 x 6 mm PDMS molds, were cultured for 14 days prior to application of compressive strain. Immediately following 24 hours compression, viscoelasticity of MONHA-encapsulated hydrogels (N = 6 per group) was measured using a Kinexus rheometer (Malvern Panalytical, Westborough, MA, USA) with an 8 mm diameter parallel plate. Rheometry measurements were performed as previously described [22]. Briefly, an 8 mm geometry was lowered onto the hydrogels to establish a calibration normal force of 0.02 N. Subsequently, an oscillatory shear-strain sweep (0.1–60%, 1.0 Hz, 25 °C) was performed to determine storage modulus (G’) and loss modulus (G”) values within the linear viscoelastic region. The storage modulus for each sample was then converted to elastic modulus (E) using the equation E = 2 * (1 + v) * G’, assuming a Poisson’s ratio (v) of 0.5 for the ECM hydrogels.

### 4.12 RNA sequencing and data sharing

The 3 mm core and the remaining periphery of MONHA-encapsulated hydrogels were isolated after compression for bulk RNA-sequencing. Three samples per strain group (i.e., 0%, 3%, 10%) at each time point (i.e., 4h, 24h) were used. Briefly, each sample was generated by pooling corresponding microregions (i.e., core or periphery) from two independent MONHA-encapsulated hydrogels, and then homogenized for 2 minutes using a Fisherbrand™ 150 Handheld Homogenizer. Total RNA was extracted using Trizol reagent (Invitrogen, 15596026, Carlsbad, CA, United States) according to the manufacturer’s protocol. 250 ng of extracted RNA was used as input to the Illumina Stranded mRNA Library prep kit as per the manufacturer’s instructions. Library quality and quantity was assessed with Agilent 2100 Bioanalyzer (Agilent Technologies, Palo Alto, CA, United States). 36 libraries were pooled together for sequencing on an Illumina NextSeq 2000 next-generation sequencing instrument with paired end 2×100bp. High-quality clean reads were filtered into FASTQ format. RNA sequencing data is deposited and accessible through NIH Gene Expression Omnibus (GEO) DataSets GSE300233.

### 4.13 Bioinformatic data analysis

RNA-seq data were analysed using the online platform Illumina Partek-Flow (version 12.6.1) (Chesterfield, Missouri, USA). Briefly, FASTQ files were subjected to sequence quality checks using “QC for Sequencing Reads” tool. Using the STAR 2.7.8a tool the reads were mapped against the mouse reference genome mm39. Aligned reads were then quantified to annotation model of mouse genome. Principal component analysis (PCA) of the top 2000 genes with greatest variance in expression was applied to all samples to delineate clustered groups between biomechanical strain durations and intensities. Differentially expressed genes (DEGs) across time (i.e., 24h vs 4h) and biomechanical strain magnitude (i.e., 0% vs 3% vs 10%) as cofactors were determined using ANOVA. Separately, pairwise comparisons between biomechanical strain intensities (i.e., 3% vs 0%, and 10% vs 0%) at each timepoint (i.e., 4h or 24h) within each hydrogel region (i.e., core or periphery) were identified using DESeq2 tool. For both analyses, the genes and transcripts with FDR corrected p-value ≤ 0.05 and |Log2 fold change| ≥ 1.50 were considered differentially expressed. Gene ontology (GO) pertaining to biological process, cellular component, and molecular function for each individual comparison was determined based on the over representation analysis (ORA) using the publicly available enrichment tool WEB-based GEne SeT AnaLysis Toolkit (https://www.webgestalt.org) [70] with FDR corrected p-value ≤ 0.05. Gene Set Enrichment Analysis (GSEA) against KEGG/Reactome databases was used to delineate altered molecular pathways across time amongst all groups and significance was set to FDR corrected p-value ≤ 0.05 and |Log2 fold change| ≥ 1.50.

### 4.14 Statistical analyses

Statistical analyses were performed using GraphPad Prism software v10.0.2 (GraphPad Software, La Jolla, CA, USA). Experimental results are illustrated using multivariable graphs and histograms, and sample sizes are specified in each figure caption. Group comparisons were determined by one-way (ANOVA) with Tukey’s multiple comparisons *post hoc* as appropriate and Cohen’s effect size d as described in **Methods 2.6** [67]. All data are shown with mean ± SD. The level of significance was set to p < 0.05.

## Acknowledgments

We thank the Neuroscience Microscopy Core at Upstate Medical University and the Blatt BioImaging Center at Syracuse University for imaging support. We thank the laboratory of Dr. Daniel J. Kelly at Trinity Centre for Biomedical Engineering, University of Dublin for their guidance and support in establishing the bioreactor system in our laboratory.

## Author contributions

A.N.S., S.P, M-T. H. T., A.K., S.G., M.P.G., A.V., A.M.B, M.V., Y.L., A.E.P., S.H. and P.S.G. designed all experiments, collected, analyzed, and interpreted the data. A.N.S and P.S.G. wrote the manuscript. All authors commented on and approved the final manuscript. S.H. and P.S.G conceived and supervised the research.

## Data and materials availability

All data needed to evaluate the conclusions in the paper are present in the paper and/or the Supplementary Materials, and RNAsequencing Data available on NIH GEO DataSets GSE300233.. Additional data related to this paper may be requested from the authors.

## Abbreviations

ATP: adenosine triphosphate
ATR: ataxia telangiectasia and Rad3 related
CNS: central nervous system
ECM: extracellular matrix
EGF: epidermal growth factor
ETC: electron transport chain
F-actin: filamentous actin
FBS: fetal bovine serum
GFAP: glial fibrillary acidic protein
HIF-1α: hypoxia-inducible factor 1 α
IL6: interleukin 6
IL17: interleukin 17
IOP: intraocular pressure
MONHA: mouse optic nerve head astrocyte
ONH: optic nerve head
PI3K-Akt: phosphoinositide 3-kinase Ak strain transforming
TGFβ2: transforming growth factor beta 2

